# Assessing substrate scope of the cyclodehydratase LynD by mRNA display-enabled machine learning models

**DOI:** 10.1101/2024.10.14.618330

**Authors:** Emma G. Steude, Henry Dieckhaus, Jarrett M. Pelton, Brian Kuhlman, Albert A. Bowers

**Affiliations:** Division of Chemical Biology and Medicinal Chemistry, UNC Eshelman School of Pharmacy, University of North Carolina at Chapel Hill, Chapel Hill, North Carolina 27599, USA; Center for Integrative Chemical Biology and Drug Discovery, Chemical Biology and Medicinal Chemistry, Eshelman School of Pharmacy, University of North Carolina at Chapel Hill, Chapel Hill, NC, 27599, USA; Department of Biochemistry and Biophysics, School of Medicine, University of North Carolina at Chapel Hill, Chapel Hill, North Carolina 27599, USA; Department of Chemistry, University of North Carolina at Chapel Hill, Chapel Hill, North Carolina 27599, USA; Lineberger Comprehensive Cancer Center, University of North Carolina at Chapel Hill, Chapel Hill, North Carolina 27599, USA

## Abstract

Many of the biosynthetic pathways for ribosomal synthesized and post-translationally modified peptide (RiPP) natural products make use of multi-domain enzymes with separate recruitment and catalysis domains that separately bind and modify peptide substrates. This “division of labor” allows RiPP enzymes to use relatively open and promiscuous active sites to perform chemistry at multiple residues within a peptide substrate seemingly regardless of the surrounding context. Defining, measuring, and predicting the seemingly broad substrate promiscuity of RiPPs necessitates high throughput assays, capable of assessing activity against very large libraries of peptides. Using mRNA display, a high throughput peptide display technology, we examine the substrate promiscuity of the RiPP cyclodehydratase, LynD. The vast substrate profiling that can be done with mRNA display enables the construction of deep learning models for accurate prediction of substrate processing by LynD. These models further inform on epistatic interactions involved in enzymatic processing. This work will facilitate the further elucidation of other RiPP enzymes and enable their use in the modification of mRNA display libraries for selection of modified peptide-based inhibitors and therapeutics.

## Introduction

Ribosomally-synthesized and post-translationally modified peptides (RiPPs) are an exciting family of natural products that have seen a surge in research due to exceptional versatility of their biosynthetic enzymes.^1^ Recent studies have shown that, in many cases, RiPP modules can be bisected into two discrete domains: a RiPP recognition element (RRE) which recognizes a leader peptide (LP) sequence, and the catalytic domain which modifies the core peptide (CP) comprising the residues in the final product.^2–4^ This decoupling of binding affinity from the catalytic site allows remarkable substrate promiscuity. For a given RiPP enzyme, if the substrate’s LP is kept constant, the active site can process multiple positions and is often tolerant of many mutations, including non-canonical amino acids, within the CP. RiPP enzymes may also be found with a variety of different active sites that carry out diverse post-translational modifications, including side chain modifications, backbone modifications, and a variety of cyclization strategies. This wide array of chemistries makes RiPP enzymes attractive tools for the elaboration of peptide-based inhibitors and therapeutics.

Recently, RIPP enzymes have been paired with mRNA display, a high throughput drug discovery tool that employs *in vitro* translation (IVT) to select for peptide binders against therapeutic targets.^5,6^ The high substrate tolerance and versatility of RIPP enzymes makes them a promising route to incorporate structural diversity into mRNA display peptide libraries.^7^ To further explore the utility of RIPP enzymes, we and others have exploited mRNA display to more extensively quantify their substrate scope and mechanism. Such efforts involve subjecting mRNA display peptide substrate libraries to enzymatic modification, then separating the enzymatically modified from unmodified sequences. The large libraries that can be generated by mRNA display allow the simultaneous testing of many millions of substrates, and analysis by next generation DNA sequencing (NGS) can enable very extensive cataloging of favorable and unfavorable enzyme substrates. Our lab has used this strategy to delineate substrate scope and requirements of the RiPP enzyme, PaaA,^8^ as well as the peptide macrocyclase activities of microbial transglutaminase (mTGase)^9^ and tyrosinase.^10^ More recently, Vinogradov et al. used mRNA display in combination with deep learning to characterize the substrate scope of LazDEF, an interesting split YcaO from the biosynthesis of lactazole, a thiopeptide (**Figure 1e**).^11,12^ Importantly, the insight gained from such substrate display studies has enabled the use of these RiPP enzymes for strategic applications in mRNA display campaigns against therapeutic targets of interest.^7,13,14^

**Figure 1.**
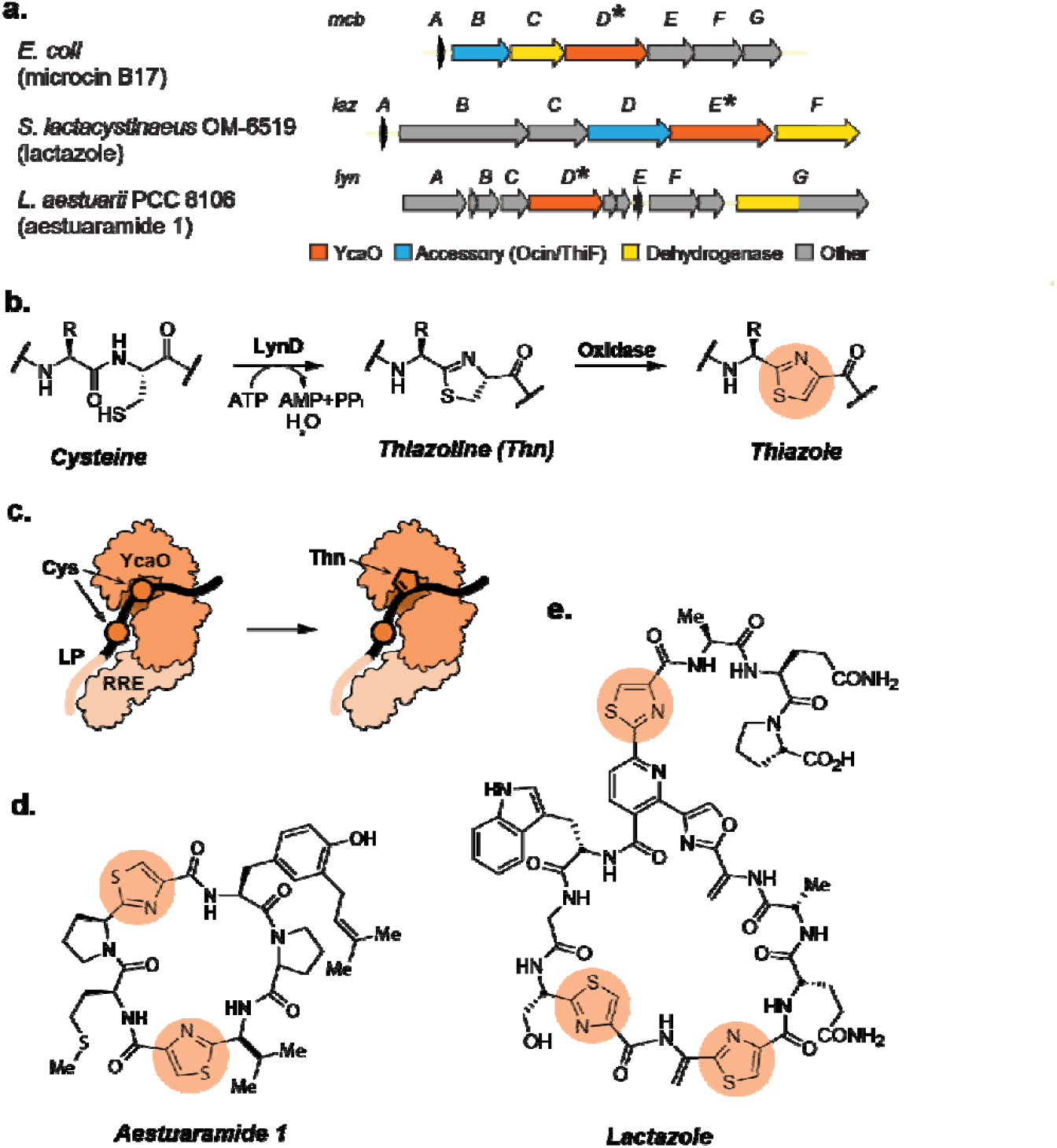
Overview of YcaO-based cyclodehydratases. (a) Gene clusters for microcin B17, lactazole, and aestuaramide 1 natural products. (b) A biosynthetic pathway from cysteine to thiazole. LynD modifies cysteines into thiazolines. Subsequent dehydrogenation by an azoline oxidase generates the final thiazole heterocycle (orange). (c) Cartoon depiction of a LynD monomer and substrate. The substrate’s leader peptide (LP) binds the RiPP recognition element (RRE) separately from the YcaO catalytic site. Only the farther downstream cysteine (Cys) reaches the catalytic site of LynD to be modified to thiazoline (Thn). (d) Chemical structures of aestuaramide 1 and lactazole natural products with thiazoles highlighted (orange).

YcaO-based cyclodehydratases are a particularly versatile family of RiPP enzymes that can carry out multiple valuable chemistries on peptide substrates.^15–19^ The first RiPP YcaO, part of the McbBCD complex involved in Microcin B17 biosynthesis, was investigated in the labs of Walsh and Kolter nearly two decades ago and shown to affect the cyclodehydrations of Cys/Ser/Thr to the corresponding thiazolines/oxazolines (**Figure 1b**).^20^ Subsequent work has shown that, within the Microcin B17 biosynthetic gene cluster (**Figure 1a**), McbD is a YcaO that catalyzes the cyclodehydration in an ATP-dependent manner and McbB is a E1-like partner protein that cooperates to form the RRE fold that recruits the substrate LP.^21–27^ Variations on this protein architecture can be found in many other YcaO-containing RiPP pathways, including thiopeptides, where the RRE can be found split from or directly fused to the YcaO, sometimes requiring the presence of additional partner proteins, such as the Ocin-ThiF-like proteins.^15,28–30^ The resulting azolines can then be converted to azoles in a final oxidation step by dehydrogenases, often also found in the clusters. In LazDEF’s case, LazDE is predicted to be a split cyclodehydratase, where LazD is an Ocin-ThiF-like protein containing the RRE and LazE is the catalytic YcaO domain that converts Cys and Ser into the respective thiazolines and oxazolines; LazF is the dehydrogenase that effects the final oxidation of these heterocycles.^12,31–34^ From a chemical standpoint, these azoles are attractive additions to peptide therapeutics because they can act as amide isosteres that add structural rigidity to RiPP natural products while reducing the hydrogen bond donating capability of the backbone amine, aiding in activity, metabolic stability, and cell uptake. Deeper understanding of their substrate scope would greatly enhance their use in the production and modification of peptide therapeutics.

Herein, we deploy substrate display in combination with deep learning to assess the substrate scope of LynD (UniProtKB A0YXD2) from aesturamide biosynthesis (**Figure 1d**). Like LazDE, LynD is a YcaO-based cyclodehydratase that is capable of converting Cys-residues to thiazolines (**Figure 1c**), but unlike LazDE, LynD does not require the assistance of an Ocin-ThiF-like partner protein (such as LazD) and has generally been observed to avoid Ser/Thr dehydration.^35^ Thus, as an enzyme from the same broad class as LazDE, but with significant differences, we expected that the study of LynD would allow us to draw meaningful comparisons in terms of substrate scope and possibly mechanism. Similar to Vinogradov et al.’s approach, we applied machine learning to significantly extend the insights from the mRNA display experiments with what proved to be very accurate predictions.^11^ Additionally, the availability of multiple crystal structures of LynD and substantial biochemical characterization already in hand for LynD and close homologs could enable a more physical interpretation of the display data than was possible in the case of LazDE.^36^ Lastly, we planned to employ a substrate display library in which the position of a constant Cys-residue was varied in relation to the LP motif. We reasoned that this library design would enable the identification of changes in substrate preferences that may result from greater degrees of freedom as the Cys-residue is further separated from the relatively static, RRE-bound LP-region. Overall, this machine learning approach reveals key structure activity relationship information for LynD that will support LynD’s further application within mRNA display, as well as the means for understanding and use of other potential YcaO-based cyclodehydratases to help produce more diverse peptide libraries.

## Results

### Streptavidin Capture Assay to Probe LynD Activity in mRNA Display

We first adapted the LazDE substrate display approach to test the substrate tolerance of LynD.^11^ In brief, this strategy uses selective alkylation/biotinylation of free cysteines in an mRNA display library to differentiate between modified and unmodified peptides in the library (**Figure 2a**). In our protocol, LynD is added to the library after the Puromycin-linked mRNA library has undergone *in vitro translation* (IVT), ribosome dissociation, and reverse transcription (RT) to pair the RNA tag with its cDNA complement. At this point, LynD will presumably modify some library members more efficiently than other members depending on their structural complementarity to the catalytic domain and ability to correctly present Cys-side chains to the active site. Subsequent treatment with biotin iodoacetamide (BIAA) will result in irreversible biotinylation of Cys-residues in peptides that have not undergone conversion to thiazolines. These unmodified sequences can then be separated from modified sequences by capture on magnetic streptavidin (SA) beads. Robustness and selectivity of this peptide modification protocol in our hands was assessed by qPCR with model substrates (**Supplementary Figure S1**) and was shown to have minimal effect on resulting cDNA concentrations, indicating that LynD-mediated thiazoline formaion is fully compatible with mRNA display methodology.

**Figure 2.**
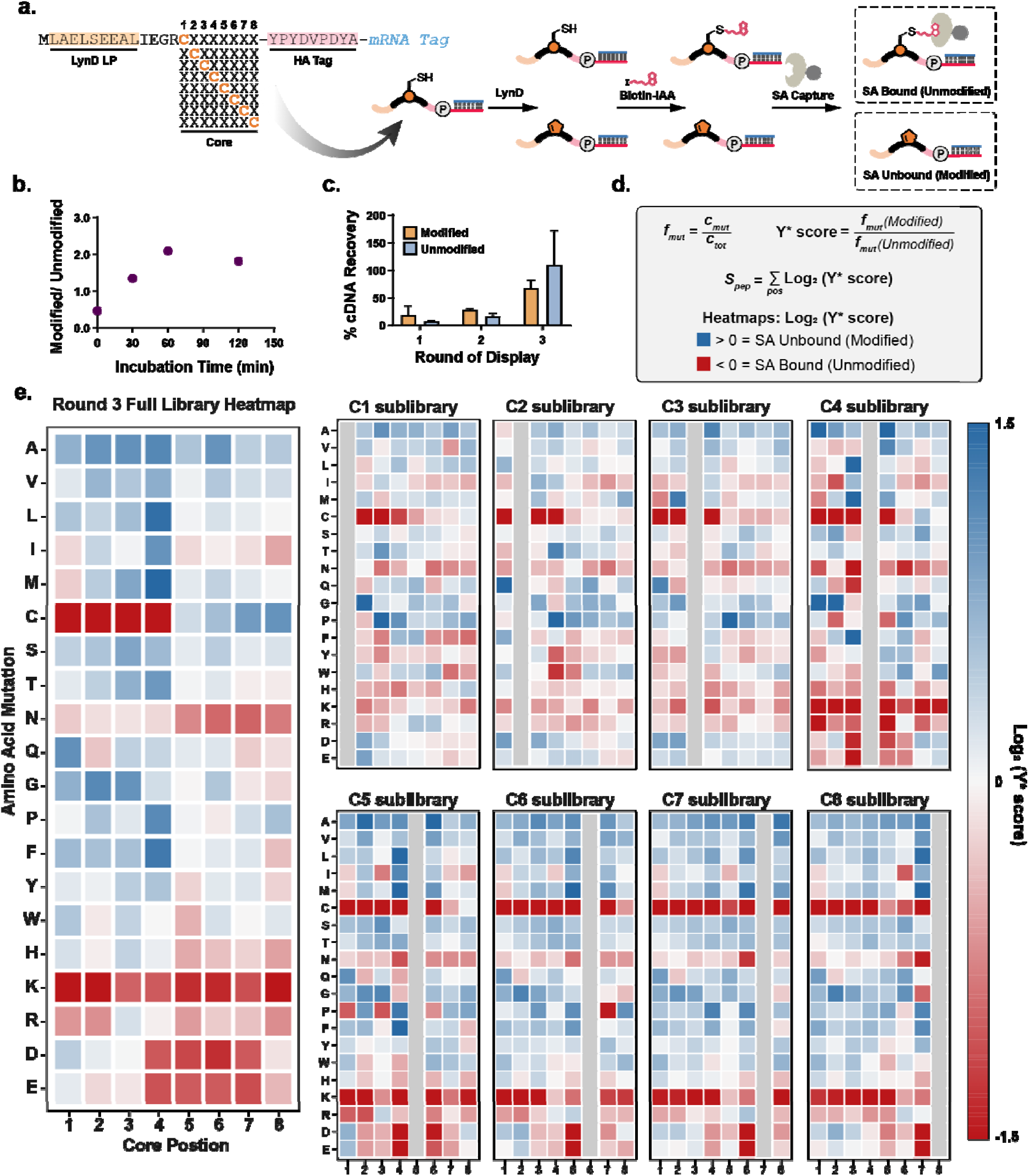
LynD streptavidin capture assay. (a) Library design and cartoon overview of LynD streptavidin capture assay. (b) LynD catalysis productivity with different incubation times. The cDNA from the modified and unmodified fract ons was quantified with qPCR to calculate the modified/ unmodified ratios. (c) Percent cDNA recovery across the three rounds of display, separated by modified and unmodified fractions. (d) Calculations for heatmaps and S-scores. (e) Heatmaps of round 3 NGS results, averaged and separated by fixed Cys position sublibraries. The horizontal axis is the core sequence position around the fixed cysteine, whereas the vertical axis shows one-letter codes of the twenty canonical amino acids that may be incorporated in the NNK library.

With a robust protocol in hand, we next turned to evaluate the promiscuity of LynD on a randomized library of substrates. For our library design, we included the experimentally defined substrate recognition LynD LP at the N-terminus, followed by a 4-residue Factor XA cleavage site, an 8-residue variable region, and a C-terminal hemagglutinin affinity (HA) purification tag. Based on a rudimentary examination of the LynD crystal structure, we anticipated the Factor XA site might provide sufficient distance between the LP and variable region (or CP) so that the CP could access the active site.^35^ Additionally, although not taken advantage of in this study, we envisioned that the site could be used to cleave the LP from modified substrates in future applications. In our design, the variable region is, in fact, a combination of 8 sublibraries with a fixed cysteine residue screened across each core position for a total theoretical diversity of roughly 1 × 10^10^ (**Figure 2a**). This library design ensures at least one Cys-residue is present in each peptide and allows us to explore the impact of the length of peptide spacing between the LP and Cys-residue undergoing modification.

We next carried out a short time course on this library to identify conditions that would leave roughly equivalent amounts of modified and unmodified library. This half-maximal concentration strategy allows for differentiation of substrates with high fitness for LynD catalysis versus those with poor fitness, both of which would be needed to train our ML models. At a concentration of roughly 30 *μ*M, LynD reached maximum activity near one hour, with a ratio of modified to unmodified of approximately 2.0 (**Figure 2b**); at 30 minutes, this ratio was reduced to near 1.0. We thus adopted the shorter reaction time for subsequent studies. As previous reports with LazDE had indicated that multiple rounds of substrate display enrichment yield better ML model accuracy, we carried out three rounds of substrate display (**Figure 2c**). For each round, we separately amplified and brought forward the SA bound (unmodified, antiselection) and unbound (modified, selection) to generate the two necessary pools for ML modeling. The cDNA was also amplified and submitted for next generation sequencing (NGS).

### Sequencing Analysis to Discern Structure-Activity Relationships for LynD Substrates

After 3 rounds of selection, over 100,000 distinct peptide sequences were returned by NGS for the selection and antiselections, combined. We initially analyzed the substrate promiscuity of LynD at the amino acid level by calculating Y* scores for a given position (**Figure 2d**).^11^ Y* scores are calculated as the ratio of frequency of mutation (*f*_*mut*_) in the modified (selection) versus unmodified (antiselection); log_2_(Y* score) values were calculated to normalize the Y* scores around zero. Positive log_2_(Y* Score) values correspond to better substrates, whereas negative values correspond to worse substrates for LynD. These values are plotted as heatmaps of amino acid mutation vs core position for the library as a whole, as well as for the eight Cys-scanning sublibraries (**Figure 2e**). The heatmaps can be used to discern LynD promiscuity at the amino acid level by highlighting positions that are more or less amenable to specific amino acid replacements.

Several substrate trends are immediately apparent from these heatmaps. First and perhaps most profoundly, cysteines are severely disfavored in core positions 1-4 relative to positions 5-8. This pattern can also be seen in the raw sequence counts for the individual sublibraries in the selection and antiselection pools (**Supplementary Figure S2**). This trend indicates that Cys-residues positioned too close to the LP recognition sequence may be prevented from undergoing enzymatic modification. Secondly, charged residues (Arg, Lys, Glu, and Asp) are broadly disfavored throughout the peptide substrates. Lys, in particular, appears to have a strong negative impact on modification, regardless of its location in the substrate, whereas Asp and Glu are most deleterious when positioned directly adjacent to the Cys-residue undergoing modification. Additionally, Cys itself is poorly tolerated when located directly next to the Cys being modified, suggesting that the formation of conjugated bis-thiazolines may be a challenge for LynD. In contrast, aliphatic amino acids are generally well tolerated by the enzyme. Ultimately, these general trends in SAR may be useful to inform future applications of LynD, but we anticipated that a more rigorous understanding of the factors controlling substrate processing might be possible by using our data to train more accurate computational models.

### Predictive Models for LynD Substrate Scope

We sought to analyze fitness at the substrate level by means of statistical fitness score, or S-score (*S*_*pep*_), which is the sum of the Log_2_(Y* Score) values across all variable positions for a specific peptide (**Figure 2d**). S-scores appeared to have good predictive power for the LazDEF system, although they have performed substantially less well in other enzyme systems that have been studied. S-scores could be calculated for each individual peptide from both the modified and unmodified fractions and then graphed by abundance (**Figure 3a**). For LynD, while there is some overlap between the fractions, the population of successfully modified peptides has a higher average S-score than the population of unmodified peptides, suggesting that S-score values alone may be able to predict whether an amino acid sequence will be modified by LynD. In contrast to the LazDEF model data, two subpopulations may be seen in the unmodified S-score plot, which may be indicative of two potential factors involved in modification ability: the substrate residues surrounding the cysteine to be modified and the cysteine’s structural distance from the LynD LP. Additionally, it should be noted that, as in the LazDE case, a majority of the naïve library S-scores fall in the range of -3 to 2, where predictions can be less accurate.

**Figure 3.**
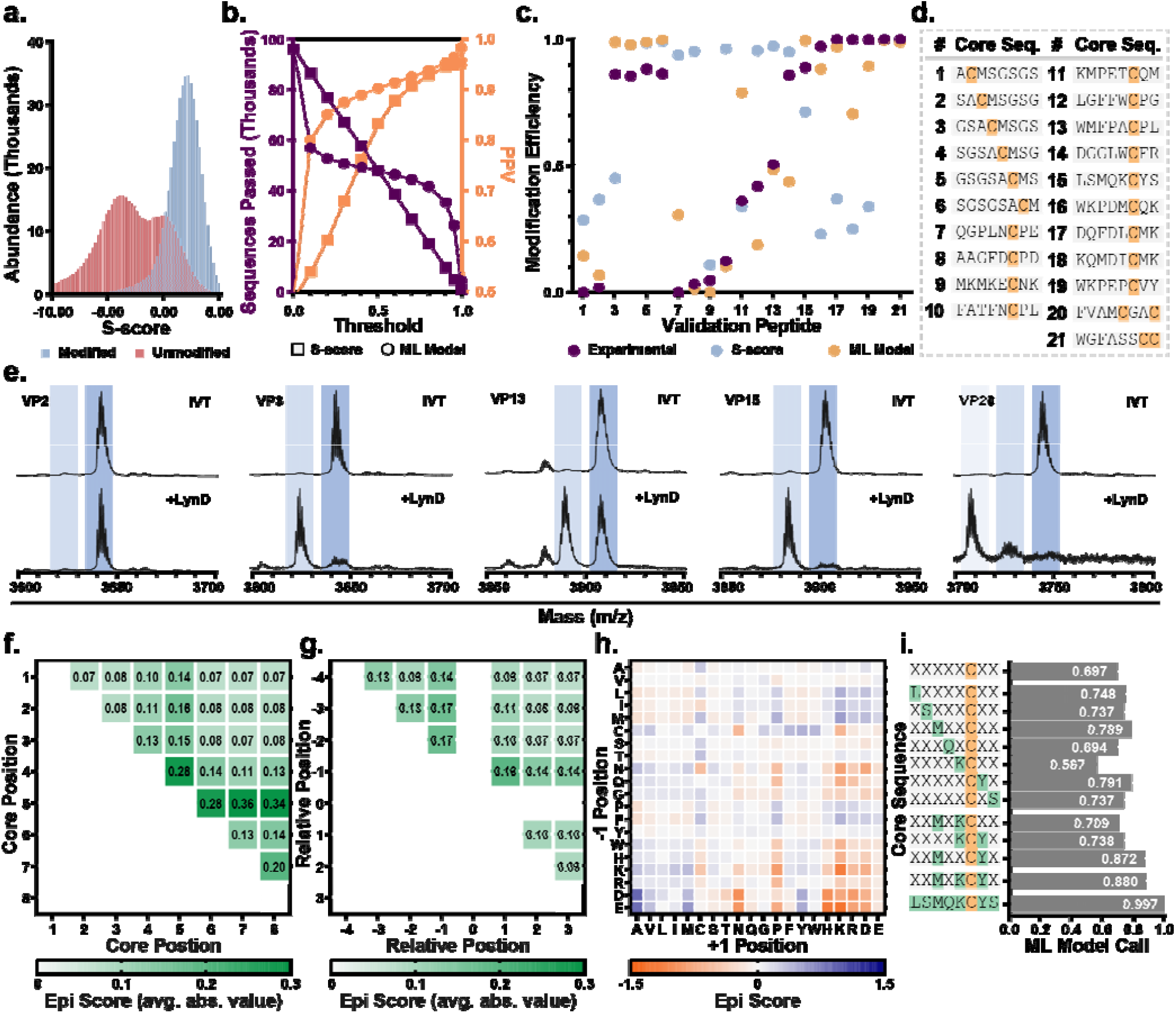
S-score and ML model comparison. (a) Abundance of peptides with each S-score, separated by modified (blue) and unmodified (pink) fractions. (b) S-score (squares) and ML model (circles) predictive model accuracy. The number of sequences past a threshold S-score value (purple) and positive predictive value (PPV, orange) are plotted according to threshold S-score value. (c) Experimentally determined (purple), S-score predicted (blue), and ML model redicted (orange) modification efficiency for validation peptides, **Genes 1-21**. (d) Core amino acid sequences for validation peptides. (e) MALDI-MS spectra for select validation peptides pre- and post-LynD treatment. The unmodified peptide mass and expected mass decrease (18 amu) for each thiazoline incorporation is highlighted with a blue band, with the lighter band representing the modified peptide mass. **Gene 20** has highlighted expected masses corresponding to modifications of either one or both cysteines. (Full set: **Supplementary Figure S4**) (f) Average positional epistas s scores for the full library. (g) Average epistasis scores for sublibraries 5-8, overlayed by relative core position (Separated by sublibrary: **Supplementary Figure S5**). (h) Epistasis between amino acids in the +1 and -1 relative positions for sublibraries 5-7. (i) ML model epistasis breakdown for validation peptide, **Gene 15**. The ML model call is shown for each core sequence, with X representing a variable amino acid position.

To improve on the accuracy of the S-score model and potentially elucidate deeper SAR trends, we next sought to train a machine learning model to the sequencing data. From the full sequencing data, the variable region of each sequence was extracted, and represented as a “one-hot” encoding. We employed a multilayer perceptron (MLP) model that was trained to predict modifiable vs unmodifiable sequences using a binary cross-entropy loss function. In agreement with the LazDEF study, we found that model architecture and sequence encoding had little effect on overall model accuracy. To evaluate the efficacy of this model and the S-score based predictions, 10% of the sequencing results were randomly assigned as a hold-out set and scored by both methods. The sequences in the hold-out set with scores above a range of threshold values were considered. For those peptide sequences, the percentage of successfully modified peptides was calculated as the positive predictive value (PPV) for each model (**Figure 3b**). The MLP model provided a higher PPV at any threshold compared to the S-score model, and the sequences passed by the MLP model followed a roughly sigmoidal trend. We also calculated the balanced accuracy (BA) of each model to evaluate predictions on both successfully and unsuccessfully modified peptides, finding that the MLP model had a BA of 92%, significantly better than the S-score model BA of 85% (**Supplementary Figure S3**). Also consistent with the LazDEF study, we found that our MLP model accuracy increased significantly with additional selection rounds, with an initial BA of 71% when trained and evaluated on the round 1 results, compared to 92% on round 3.

To test the predictive capability of both the S-score and MLP methods, we compared predicted modification efficiencies from both models to experimentally determined values for select peptides (**Figure 3c,d**). We chose 15 peptides that exhibited a range of predicted modification efficiencies, including several instances where the S-score and MLP model exhibited significant differences in their predictions. Additionally, we included another 6 peptides for validation that sampled different distances between the Cys-residue and LP to confirm the observed sublibrary preference. These genes were translated *in vitro* and incubated with LynD for 30 minutes as in the display protocol. Modification efficiency was then quantified by MALDI-MS (**Figure 3e**).

In general, the results showed that the MLP model outperformed S-score. First, we evaluated Cys location dependance. **Genes 1-6** were designed to screen the cysteine at each core position, using GS linkers of different lengths to vary the distance away from the LynD leader peptide. Indeed, only **Genes 3-6**, with Cys-residues beyond core position 3, are modified by LynD in this assay. This strongly suggests that we are able to control the site of modification by LynD in an mRNA context by altering the position of the Cys-residue relative to the LP. In regions sufficiently removed from the LP, both S-score and MLP model score can be used to predict modification efficiency with reasonable accuracy. To better compare the S-score and MLP model accuracy, we designed **Genes 7-19** to include a range of predictive values with contrasting model scores, all with a fixed Cys at core position 6. For these test genes, the experimental values were closer to the MLP-score than the S-score value in all but two cases: **Genes 11** and **14** had MLP predictive values in the middle range of scores, but in one case MLP over-predicted modification and in another it under-predicted. It is not unexpected that MLP model predictions might deviate somewhat with midrange predictions that are less decisive. The maximum difference in experimental LynD modification efficiency compared to our MLP-score predictions (Δ_max_) is 0.43, comparable to the LazDE model, which exhibited a Δ_max_ of 0.36. Notably, several peptides with multiple cysteines were also tested in these assays. **Genes 20** and **21** display two cysteines in the core region, as separated and adjacent residues respectively. Both genes exhibited double modification after the 30-minute incubation, in good agreement with the high predictive values from both models. This result is also somewhat to be expected, as the natural substrate of LynD includes multiple cysteines in its core region, so LynD is known to be capable of performing iterative cyclizations.

### Evaluating Epistatic Contributions to Substrate Fitness

Finally, given the robust performance of our MLP model and good correlation with experimental results, we sought to use the model to examine higher order, epistatic effects that could potentially have led to deviations from S-score predictions in certain substrates. To this end, we employed the model to calculate epistasis (*epi*) scores for each randomized position in the fixed cysteine library. Positive *epi* scores correspond to synergistic effects between amino acids at two given positions, while negative scores indicate deleterious effects. Average absolute *epi* scores can be used to probe overall coupling between positions in the library as a whole, with values farther from zero corresponding to higher coupling (**Figure 3f**). Given the complexity of our library, the map of average absolute *epi* scores is dominated by combinations of interactions near the fixed Cys-residues in the underlying C-terminal sublibraries. Thus, overall, the epistasis score between core positions 5 and 7 appear the highest, at 0.36.

We then extracted and overlayed the *epi* scores for sublibraries 5-8, which undergo the most extensive modification, to gain better insights. This analysis shows that these sublibraries exhibit the highest epistatic coupling between the -1 and +1 positions relative to the cysteine, at 0.19 (**Figure 3g**), and much subtler couplings at other positions. This pattern is consistent with data on the LazDE system, as well as several other known and characterized YcaO-based cyclodehydratases. The coupling between specific amino acids in these -1 and +1 positions can also be extracted and shown in a more detailed heatmap (**Figure 3h**). Many amino acid pairs show little epistasis, with values near zero, but some pairs exhibit stronger coupling, either enhancing or diminishing modification, sometimes in unexpected ways. For example, the *epi* score is strongly negative when Lys and Pro are in positions -1 and +1 respectively. This suggests that, as expected, although these two residues decrease fitness on their own, they can do so to an even greater extent when found in conjunction. In contrast, when Ala is present in the +1 position it can help overcome the effects of Asp in the -1 position, resulting in a relatively larger positive *epi* score. Similarly, more extensive interactions can also be found to help save other disfavored substrates. In **Gene 15**, Met at the -3 position combined with Tyr in the +1 position can offset the penalty of Lys in the -1 position that is broadly observed in the library (**Figure 3i**). Still other combinations can help to allow simultaneous modification of multiple Cys-residues (**Supplementary Figure S6**). Overall, the results from the MLP model with the randomized library yielded epistasis information about how amino acids in conversation with one another may affect the success of LynD modification.

## Discussion

Herein we have applied a machine learning guided substrate display assay to assess the substrate scope of the YcaO-based cyclodehydratase, LynD. The substrate scope bares significant similarities and differences to that of LazDE. Both LynD and LazDE exhibit strong dependence on residues in the -1 and +1 positions relative to the Cys undergoing modification. In both sets of sequencing data, these residues witness the greatest variance in log_2_(Y* Score) values and the most significant epistatic coupling as inferred from *epi* scores. The enzymes also exhibit some level of similarity in their amino acid preferences at these and other positions. In general, neither enzyme appears to handle formation of multiple, adjacent thiazoles well, resulting in reduction of Cys-residues in the modified selection pool and enrichment in the antiselections. Charged residues also appear to negatively impact both enzymes, although to different extents: while LazDE disfavors all charged residues, it is much more sensitive to negative charge (Asp and Glu) than positive (Lys and Arg). LynD also disfavors all charged residues, but is more sensitive to positive charge, especially Lys, than negative charge. Lastly, LazDE strongly prefers small hydrophobics (Ala and Gly) and disfavors aromatics at the -1 and +1 positions, while LynD appears more promiscuous in the hydrophobic space. The latter difference in promiscuity is also partly reflected in the higher average MLP-score for LynD versus LazDE. In the long term, similar experimental and modelling pipelines against other enzyme family members may help to identify differences in structure-activity relationships with greater resolution.

The ready availability of the crystal structure of LynD provides additional context to our substrate display results.^35^ Importantly, the LynD structure shows that LynD exists as an intertwined dimer, with the LP recognition sequence bound roughly 14 Å from the ATP in the active site of the enzyme (**Supplementary Figure S8**). Depending on the degree of protein dynamics, this distance could vary significantly during the process of catalysis, but the requisite binding of the LP should still put a theoretical limit on the allowable tether between the LP and the Cys-residue undergoing modification. Such a limit is in good agreement with our selection data, which shows a stark preference for substrates bearing Cys-residues at positions 5-8. Peptides with a Cys-residue at position 4 show intermediate levels of modification, consistent with there being a degree of flexibility in the protein:peptide interaction that permits some transient access of the residue to the catalytic domain. Given that LazDE requires the involvement of the LazD Ocin-ThiF-like protein and its own, presumed protein:protein interactions with LazE, it is unclear what kind of distance constraints this enzyme complex would exhibit and what level of fidelity can be levied over the position of modification through the length of the LP tether.

Additionally, the electrostatic potential of the LynD active site as inferred from the crystal structure could help to explain why the enzyme generally disfavors charged residues near the Cys undergoing modification. Using a docked structure of the native substrate of LynD as well as a similar docking to an AlphaFold 3 (AF3)-generated structure of LazE (**Supplementary Figure S9**), it is possible to estimate the average electrostatic potential of regions directly surrounding the -3, -2, -1, 0, +1, +2, and +3 positions of the peptide substrate. While this is only an approximation for heuristic purposes, the high homology within the YcaO family members lends itself to a high confidence AF3 prediction for LazE. Nevertheless, the active site electrostatic potential for LazE is predicted to be significantly more negative than for LynD, potentially causing more deleterious interactions with negatively charged Asp and Glu-residues in LazE substrates. In contrast, the more neutral electrostatic potential of LynD may hinder its activity with positively charged Lys-residues. While speculative, these insights may lend themselves to the nomination of alternative YcaO-based cyclodehydratases for applications against charged substrates or even the identification of still more promiscuous members of the family.

Cumulatively, this work provides substantial new insight into the substrate scope of LynD and a validated model for high confidence prediction of additional activity. We anticipate that these data and tools can be used to harness LynD for new applications, particularly in the modification of mRNA display libraries for selection against therapeutic targets. The Cys-dehydrations carried out by LynD and related family members are high value modifications for peptide inhibitors because the dehydration creates a rigidified amide bond isostere, while also removing a hydrogen-bond donor from the peptide backbone.^37–40^ Being able to control the placement of this modification through altering its proximity to the leader peptide could significantly benefit LynD’s applications in libraries and allow synergistic modifications with other Cys-targeting chemistries and chemical transformations. Additionally, we expect that this work will enable similar studies against other family members, eventually yielding greater insight into the enzyme family. Machine learning is a powerful tool for the analysis of large datasets and its natural combination with mRNA display appears poised to make contributions in many facets of protein biochemistry and enzymology.

## ASSOCIATED CONTENT

The supporting information is available free of charge at *hyperlink here*

Experimental methods and supporting figures (PDF)

## Notes

The authors declare no competing financial interests.

## ACKNOWLEDGMENTS

This work was supported by grants from the NSF (CHE-2204094 to A. A. B.) and NIH to B. A. K. (R35GM131923). E.G.S. acknowledges support from the University of North Carolina at Chapel Hill Department of Chemistry’s J. Thurman Freeze Research Award. H.D. acknowledges support from the NSF (DGE-2040435 to H.D.), as well as a Pre-doctoral Fellowship from the American Foundation for Pharmaceutical Education. This work utilized the resources of the UNC Longleaf high-performance computing cluster.

## ACCESSION CODES

- LynD UniProt ID: A0YXD2

