## Supporting Information for "Assessing substrate scope of the cyclodehydratase LynD by mRNA display-enabled machine learning models"

Emma G. Steude,^1,2^ Henry Dieckhaus,^1,3^ Jarrett M. Pelton,^1,2^ Brian Kuhlman,^3,5^ Albert A. Bowers^1,2,4,5^*

1. Division of Chemical Biology and Medicinal Chemistry, UNC Eshelman School of Pharmacy, University of North Carolina at Chapel Hill, Chapel Hill , North Carolina 27599, USA
2. Center for Integrative Chemical Biology and Drug Discovery, Chemical Biology and Medicinal Chemistry, Eshelman School of Pharmacy, University of North Carolina at Chapel Hill, Chapel Hill, NC, 27599, USA
3. Department of Biochemistry and Biophysics, School of Medicine, University of North Carolina at Chapel Hill, Chapel Hill, North Carolina 27599, USA
4. Department of Chemistry, University of North Carolina at Chapel Hill, Chapel Hill, North Carolina 27599, USA
5. Lineberger Comprehensive Cancer Center, University of North Carolina at Chapel Hill, Chapel Hill, North Carolina 27599, USA

### General information

##### Reagents

Reagents for molecular biology were purchased from New England BioLabs (NEB), Thermo Fisher Scientific, and Fisher Scientific. Specifically, PURExpress KITS (E6840S – standard translations, and E6850Z – custom for mRNA display), T7 RNA Polymerase (M0251L), Q5 DNA Polymerase (M0491L) were purchased from New England Biolabs. T4 RNA ligase I (PR-M1051), RQ1 DNase (PR-M6101), and M-MLV Reverse Transcriptase, RNase H Minus, Point Mutant (PRM3683) were purchased from Promega through Fisher Scientific. Anti-HA magnetic beads (88836), HA synthetic peptide (26184), and M-280 Streptavidin Dynabeads (11205D) were purchased from Thermo Fisher Scientific. Bulk solvents were purchased from Fisher Scientific. DNA gene fragments were purchased from Twist bioscience. Primers were purchased from Integrated DNA Technologies. Next Generation Sequencing (NGS) was performed by Azenta using their Amplicon-EZ (150-500bp) service.

##### Software

Software used for data analysis comprised of Qiagen CLC Main Workbench (NGS analysis), Anaconda navigator Jupyter Notebook v6.3.0 (Python Script), Graphpad Prism v9.2.0, and Microsoft office suite. Adobe Illustrator was used to assemble figures. Finally, in house python scripts were used to analyze NGS data.

##### Definitions

For complete definitions of NGS data calculations, please see Ref. 1 below.

### Methods

#### List of Primer, Library, and Gene Sequences

Primer Sequences:

| # | Name | Purpose | DNA Sequence |
| --- | --- | --- | --- |
| P1 | Lib Fwd Amp | Fwd amplification of NNK randomized library | TAATACGACTCACTATAGGGTTAACTTTAAGAAGGAGATATACATATGCTGGCGG |
| P2 | Lib Fwd Ext | Fwd extension primer for NNK randomized library | CATATGCTGGCGGAATTATCAGAAGAGGCTTTAATTGAAG |
| P3 | Lib Rev Ext 1 | Rev extension primer for NNK randomized library with TGT codon at position 1 | CGGGAACATCATAAGGATAAGAACCAGAACCMNNMNNMNNMNNMNNMNNMNN**ACA**ACGGCCTTCAATTAAAGCCTC |
| P4 | Lib Rev Ext 2 | Rev extension primer for NNK randomized library with TGT codon at position 2 | CGGGAACATCATAAGGATAAGAACCAGAACCMNNMNNMNNMNNMNNMNN**ACA**MNNACGGCCTTCAATTAAAGCCTC |
| P5 | Lib Rev Ext 3 | Rev extension primer for NNK randomized library with TGT codon at position 3 | CGGGAACATCATAAGGATAAGAACCAGAACCMNNMNNMNNMNNMNN**ACA**MNNMNNACGGCCTTCAATTAAAGCCTC |
| P6 | Lib Rev Ext 4 | Rev extension primer for NNK randomized library with TGT codon at position 4 | CGGGAACATCATAAGGATAAGAACCAGAACCMNNMNNMNNMNN**ACA**MNNMNNMNNACGGCCTTCAATTAAAGCCTC |
| P7 | Lib Rev Ext 5 | Rev extension primer for NNK randomized library with TGT codon at position 5 | CGGGAACATCATAAGGATAAGAACCAGAACCMNNMNNMNN**ACA**MNNMNNMNNMNNACGGCCTTCAATTAAAGCCTC |
| P8 | Lib Rev Ext 6 | Rev extension primer for NNK randomized library with TGT codon at position 6 | CGGGAACATCATAAGGATAAGAACCAGAACCMNNMNN**ACA**MNNMNNMNNMNNMNNACGGCCTTCAATTAAAGCCTC |
| P9 | Lib Rev Ext 7 | Rev extension primer for NNK randomized library with TGT codon at position 7 | CGGGAACATCATAAGGATAAGAACCAGAACCMNN**ACA**MNNMNNMNNMNNMNNMNNACGGCCTTCAATTAAAGCCTC |
| P10 | Lib Rev Ext 8 | Rev extension primer for NNK randomized library with TGT codon at position 8 | CGGGAACATCATAAGGATAAGAACCAGAACC**ACA**MNNMNNMNNMNNMNNMNNMNNACGGCCTTCAATTAAAGCCTC |
| P11 | Lib Rev Amp | Rev amplification of NNK randomized library | TTTCCGCCCCCCGTCTCAACTTCCACTCCCGCTACCGGCATAATCGGGAACATCATAAG |
| P12 | NGS Adapter Fwd | Fwd primer for Amplicon EZ NGS of mRNA display libraries | ACACTCTTTCCCTACACGACGCTCTTCCGATCTTAATACGACTCACTATAGGGTTAACTTTAAG |
| P13 | NGS Adapter Rev 1 | Rev primer for Amplicon EZ NGS of mRNA display libraries | GACTGGAGTTCAGACGTGTGCTCTTCCGATCT**TGC**TTTCCGCCCCCCGTCTC |
| P14 | NGS Adapter Rev 2 | Rev primer for Amplicon EZ NGS of mRNA display libraries | GACTGGAGTTCAGACGTGTGCTCTTCCGATCT**CTG**TTTCCGCCCCCCGTCTC |
| P15 | NGS Adapter Rev 3 | Rev primer for Amplicon EZ NGS of mRNA display libraries | GACTGGAGTTCAGACGTGTGCTCTTCCGATCT**GAT**TTTCCGCCCCCCGTCTC |
| P16 | NGS Adapter Rev 4 | Rev primer for Amplicon EZ NGS of mRNA display libraries | GACTGGAGTTCAGACGTGTGCTCTTCCGATCT**GGA**TTTCCGCCCCCCGTCTC |
| P17 | T7 Prom | T7 promoter for amplifying IVT peptides | GAAATTAATACGACTCACTATAGGGG |
| P18 | T7 Term | T7 terminator for amplifying IVT peptides | GCTAGTTATTGCTCAGCGG |

Assay Validation Test Gene Sequence:

G1 – Test gene modeled from LynD’s aesturamide natural product for streptavidin pull down assay validation:

GTGGCCGGCATCACCCAGGTGCGGTTGCTGGCGCCTATATCGCCGACATCACCGATGGGGAAGATCGGGCTCGCCACTTCGGGCTCATGAGCGCTTGTTTCGTAATACGACTCACTATAGGGTTAACTTTAAGAAGGAGATATACATATGCTGGCCGAACTTTCTGAGGAAGCCCTTGGCGGAAACGGAGTAGACGCTTCAGCCTGTATGCCGTGCTATCCGTCGTATGACGGTTCGTACCCGTATGATGTGCCAGACTACGCCGGAGAGCTGGCACGTCCTTGAGACGGGGGGCGGAAA

Full Library Sequence:

L1 – NNK cysteine screen library (assembled from primers)*:

*below is indicative of TGT at position 1. In full library, TGT is scanned across all 8 randomized positions.

TAATACGACTCACTATAGGGTTAACTTTAAGAAGGAGATATACATATGCTGGCGGAATTATCAGAAGAGGCTTTAATTGAAGGCCGT**TGT**NNKNNKNNKNNKNNKNNKNNKGGTTCTGGTTCTTATCCTTATGATGTTCCCGATTATGCCGGTAGCGGGAGTGGAAGTTGAGACGGGGGGCGGAAA

Validation Peptide Gene Sequences:

General validation peptide sequence: MLAELSEEALIEGR —Core— GSGSYPYDVPDYA

| # | Core AA Sequence | Test | DNA Sequence |
| --- | --- | --- | --- |
| VP1 | ACMSGSGS | Distance | GCGAAATTAATACGACTCACTATAGGGGAATTGTGAGCGGATAACAATTCCCCTCTAGAAATAATTTTGTTTAACTTTAAGAAGGAGATATACATATGCTGGCCGAACTTAGCGAAGAAGCGTTGATCGAGGGGCGCGCATGCATGAGTGGCTCGGGCTCTGGGTCTGGGAGCTACCCTTACGATGTCCCGGATTACGCCTAATAGCGCATTGGAAGTGGATAACGGATCCGAATTCGAGCTCCGTCGACAAGCTTGCGGCCGCACTCGAGTGAGATCCGGCTGCTAACAAAGCCCGAAAGGAAGCTGAGTTGGCTGCTGCCACCGCTGAGCAATAACTAGCATAACC |
| VP2 | SACMSGSG | Distance | GCGAAATTAATACGACTCACTATAGGGGAATTGTGAGCGGATAACAATTCCCCTCTAGAAATAATTTTGTTTAACTTTAAGAAGGAGATATACATATGCTGGCCGAACTTAGCGAAGAAGCGTTGATCGAGGGGCGCTCAGCATGCATGTCAGGGTCCGGCGGGTCTGGGAGCTACCCTTACGATGTCCCGGATTACGCCTAATAGCGCATTGGAAGTGGATAACGGATCCGAATTCGAGCTCCGTCGACAAGCTTGCGGCCGCACTCGAGTGAGATCCGGCTGCTAACAAAGCCCGAAAGGAAGCTGAGTTGGCTGCTGCCACCGCTGAGCAATAACTAGCATAACC |
| VP3 | GSACMSGS | Distance | GCGAAATTAATACGACTCACTATAGGGGAATTGTGAGCGGATAACAATTCCCCTCTAGAAATAATTTTGTTTAACTTTAAGAAGGAGATATACATATGCTGGCCGAACTTAGCGAAGAAGCGTTGATCGAGGGGCGCGGGTCTGCTTGTATGTCGGGGTCAGGGTCTGGGAGCTACCCTTACGATGTCCCGGATTACGCCTAATAGCGCATTGGAAGTGGATAACGGATCCGAATTCGAGCTCCGTCGACAAGCTTGCGGCCGCACTCGAGTGAGATCCGGCTGCTAACAAAGCCCGAAAGGAAGCTGAGTTGGCTGCTGCCACCGCTGAGCAATAACTAGCATAACC |
| VP4 | SGSACMSG | Distance | GCGAAATTAATACGACTCACTATAGGGGAATTGTGAGCGGATAACAATTCCCCTCTAGAAATAATTTTGTTTAACTTTAAGAAGGAGATATACATATGCTGGCCGAACTTAGCGAAGAAGCGTTGATCGAGGGGCGCTCAGGGAGCGCGTGCATGAGCGGTGGGTCTGGGAGCTACCCTTACGATGTCCCGGATTACGCCTAATAGCGCATTGGAAGTGGATAACGGATCCGAATTCGAGCTCCGTCGACAAGCTTGCGGCCGCACTCGAGTGAGATCCGGCTGCTAACAAAGCCCGAAAGGAAGCTGAGTTGGCTGCTGCCACCGCTGAGCAATAACTAGCATAACC |
| VP5 | GSGSACMS | Distance | GCGAAATTAATACGACTCACTATAGGGGAATTGTGAGCGGATAACAATTCCCCTCTAGAAATAATTTTGTTTAACTTTAAGAAGGAGATATACATATGCTGGCCGAACTTAGCGAAGAAGCGTTGATCGAGGGGCGCGGCTCTGGCTCCGCTTGCATGTCGGGGTCTGGGAGCTACCCTTACGATGTCCCGGATTACGCCTAATAGCGCATTGGAAGTGGATAACGGATCCGAATTCGAGCTCCGTCGACAAGCTTGCGGCCGCACTCGAGTGAGATCCGGCTGCTAACAAAGCCCGAAAGGAAGCTGAGTTGGCTGCTGCCACCGCTGAGCAATAACTAGCATAACC |
| VP6 | SGSGSACM | Distance | GCGAAATTAATACGACTCACTATAGGGGAATTGTGAGCGGATAACAATTCCCCTCTAGAAATAATTTTGTTTAACTTTAAGAAGGAGATATACATATGCTGGCCGAACTTAGCGAAGAAGCGTTGATCGAGGGGCGCTCAGGCTCAGGTTCAGCTTGTATGGGGTCTGGGAGCTACCCTTACGATGTCCCGGATTACGCCTAATAGCGCATTGGAAGTGGATAACGGATCCGAATTCGAGCTCCGTCGACAAGCTTGCGGCCGCACTCGAGTGAGATCCGGCTGCTAACAAAGCCCGAAAGGAAGCTGAGTTGGCTGCTGCCACCGCTGAGCAATAACTAGCATAACC |
| VP7 | QGPLNCPE | Range of ML model scores | GCGAAATTAATACGACTCACTATAGGGGAATTGTGAGCGGATAACAATTCCCCTCTAGAAATAATTTTGTTTAACTTTAAGAAGGAGATATACATATGCTGGCCGAACTTAGCGAAGAAGCGTTGATCGAGGGGCGCCAAGGTCCGTTAAACTGCCCTGAGGGGTCTGGGAGCTACCCTTACGATGTCCCGGATTACGCCTAATAGCGCATTGGAAGTGGATAACGGATCCGAATTCGAGCTCCGTCGACAAGCTTGCGGCCGCACTCGAGTGAGATCCGGCTGCTAACAAAGCCCGAAAGGAAGCTGAGTTGGCTGCTGCCACCGCTGAGCAATAACTAGCATAACC |
| VP8 | AAGFDCPD | Range of ML model scores | GCGAAATTAATACGACTCACTATAGGGGAATTGTGAGCGGATAACAATTCCCCTCTAGAAATAATTTTGTTTAACTTTAAGAAGGAGATATACATATGCTGGCCGAACTTAGCGAAGAAGCGTTGATCGAGGGGCGCGCTGCTGGGTTTGACTGTCCTGACGGGTCTGGGAGCTACCCTTACGATGTCCCGGATTACGCCTAATAGCGCATTGGAAGTGGATAACGGATCCGAATTCGAGCTCCGTCGACAAGCTTGCGGCCGCACTCGAGTGAGATCCGGCTGCTAACAAAGCCCGAAAGGAAGCTGAGTTGGCTGCTGCCACCGCTGAGCAATAACTAGCATAACC |
| VP9 | MKMKECNK | Range of ML model scores | GCGAAATTAATACGACTCACTATAGGGGAATTGTGAGCGGATAACAATTCCCCTCTAGAAATAATTTTGTTTAACTTTAAGAAGGAGATATACATATGCTGGCCGAACTTAGCGAAGAAGCGTTGATCGAGGGGCGCATGAAAATGAAAGAATGTAATAAAGGGTCTGGGAGCTACCCTTACGATGTCCCGGATTACGCCTAATAGCGCATTGGAAGTGGATAACGGATCCGAATTCGAGCTCCGTCGACAAGCTTGCGGCCGCACTCGAGTGAGATCCGGCTGCTAACAAAGCCCGAAAGGAAGCTGAGTTGGCTGCTGCCACCGCTGAGCAATAACTAGCATAACC |
| VP10 | FATFNCPL | Range of ML model scores | GCGAAATTAATACGACTCACTATAGGGGAATTGTGAGCGGATAACAATTCCCCTCTAGAAATAATTTTGTTTAACTTTAAGAAGGAGATATACATATGCTGGCCGAACTTAGCGAAGAAGCGTTGATCGAGGGGCGCTTTGCTACTTTTAACTGTCCTCTTGGGTCTGGGAGCTACCCTTACGATGTCCCGGATTACGCCTAATAGCGCATTGGAAGTGGATAACGGATCCGAATTCGAGCTCCGTCGACAAGCTTGCGGCCGCACTCGAGTGAGATCCGGCTGCTAACAAAGCCCGAAAGGAAGCTGAGTTGGCTGCTGCCACCGCTGAGCAATAACTAGCATAACC |
| VP11 | KMPETCQM | Range of ML model scores | GCGAAATTAATACGACTCACTATAGGGGAATTGTGAGCGGATAACAATTCCCCTCTAGAAATAATTTTGTTTAACTTTAAGAAGGAGATATACATATGCTGGCCGAACTTAGCGAAGAAGCGTTGATCGAGGGGCGCAAGATGCCTGAAACCTGTCAAATGGGGTCTGGGAGCTACCCTTACGATGTCCCGGATTACGCCTAATAGCGCATTGGAAGTGGATAACGGATCCGAATTCGAGCTCCGTCGACAAGCTTGCGGCCGCACTCGAGTGAGATCCGGCTGCTAACAAAGCCCGAAAGGAAGCTGAGTTGGCTGCTGCCACCGCTGAGCAATAACTAGCATAACC |
| VP12 | LGFFWCPG | Range of ML model scores | GCGAAATTAATACGACTCACTATAGGGGAATTGTGAGCGGATAACAATTCCCCTCTAGAAATAATTTTGTTTAACTTTAAGAAGGAGATATACATATGCTGGCCGAACTTAGCGAAGAAGCGTTGATCGAGGGGCGCCTGGGGTTCTTCTGGTGTCCAGGCGGGTCTGGGAGCTACCCTTACGATGTCCCGGATTACGCCTAATAGCGCATTGGAAGTGGATAACGGATCCGAATTCGAGCTCCGTCGACAAGCTTGCGGCCGCACTCGAGTGAGATCCGGCTGCTAACAAAGCCCGAAAGGAAGCTGAGTTGGCTGCTGCCACCGCTGAGCAATAACTAGCATAACC |
| VP13 | WMFPACPL | Range of ML model scores | GCGAAATTAATACGACTCACTATAGGGGAATTGTGAGCGGATAACAATTCCCCTCTAGAAATAATTTTGTTTAACTTTAAGAAGGAGATATACATATGCTGGCCGAACTTAGCGAAGAAGCGTTGATCGAGGGGCGCTGGATGTTTCCTGCCTGCCCACTGGGGTCTGGGAGCTACCCTTACGATGTCCCGGATTACGCCTAATAGCGCATTGGAAGTGGATAACGGATCCGAATTCGAGCTCCGTCGACAAGCTTGCGGCCGCACTCGAGTGAGATCCGGCTGCTAACAAAGCCCGAAAGGAAGCTGAGTTGGCTGCTGCCACCGCTGAGCAATAACTAGCATAACC |
| VP14 | DGGLWCFR | Range of ML model scores | GCGAAATTAATACGACTCACTATAGGGGAATTGTGAGCGGATAACAATTCCCCTCTAGAAATAATTTTGTTTAACTTTAAGAAGGAGATATACATATGCTGGCCGAACTTAGCGAAGAAGCGTTGATCGAGGGGCGCGACGGGGGTCTTTGGTGTTTTCGTGGGTCTGGGAGCTACCCTTACGATGTCCCGGATTACGCCTAATAGCGCATTGGAAGTGGATAACGGATCCGAATTCGAGCTCCGTCGACAAGCTTGCGGCCGCACTCGAGTGAGATCCGGCTGCTAACAAAGCCCGAAAGGAAGCTGAGTTGGCTGCTGCCACCGCTGAGCAATAACTAGCATAACC |
| VP15 | LSMQKCYS | Range of ML model scores | GCGAAATTAATACGACTCACTATAGGGGAATTGTGAGCGGATAACAATTCCCCTCTAGAAATAATTTTGTTTAACTTTAAGAAGGAGATATACATATGCTGGCCGAACTTAGCGAAGAAGCGTTGATCGAGGGGCGCCTGTCAATGCAAAAATGCTATTCTGGGTCTGGGAGCTACCCTTACGATGTCCCGGATTACGCCTAATAGCGCATTGGAAGTGGATAACGGATCCGAATTCGAGCTCCGTCGACAAGCTTGCGGCCGCACTCGAGTGAGATCCGGCTGCTAACAAAGCCCGAAAGGAAGCTGAGTTGGCTGCTGCCACCGCTGAGCAATAACTAGCATAACC |
| VP16 | WKPDMCQK | Range of ML model scores | GCGAAATTAATACGACTCACTATAGGGGAATTGTGAGCGGATAACAATTCCCCTCTAGAAATAATTTTGTTTAACTTTAAGAAGGAGATATACATATGCTGGCCGAACTTAGCGAAGAAGCGTTGATCGAGGGGCGCTGGAAGCCAGACATGTGTCAAAAAGGGTCTGGGAGCTACCCTTACGATGTCCCGGATTACGCCTAATAGCGCATTGGAAGTGGATAACGGATCCGAATTCGAGCTCCGTCGACAAGCTTGCGGCCGCACTCGAGTGAGATCCGGCTGCTAACAAAGCCCGAAAGGAAGCTGAGTTGGCTGCTGCCACCGCTGAGCAATAACTAGCATAACC |
| VP17 | DQFDLCMK | Range of ML model scores | GCGAAATTAATACGACTCACTATAGGGGAATTGTGAGCGGATAACAATTCCCCTCTAGAAATAATTTTGTTTAACTTTAAGAAGGAGATATACATATGCTGGCCGAACTTAGCGAAGAAGCGTTGATCGAGGGGCGCGATCAATTCGATCTGTGCATGAAAGGGTCTGGGAGCTACCCTTACGATGTCCCGGATTACGCCTAATAGCGCATTGGAAGTGGATAACGGATCCGAATTCGAGCTCCGTCGACAAGCTTGCGGCCGCACTCGAGTGAGATCCGGCTGCTAACAAAGCCCGAAAGGAAGCTGAGTTGGCTGCTGCCACCGCTGAGCAATAACTAGCATAACC |
| VP18 | KQMDICMK | Range of ML model scores | GCGAAATTAATACGACTCACTATAGGGGAATTGTGAGCGGATAACAATTCCCCTCTAGAAATAATTTTGTTTAACTTTAAGAAGGAGATATACATATGCTGGCCGAACTTAGCGAAGAAGCGTTGATCGAGGGGCGCAAGCAGATGGACATTTGTATGAAAGGGTCTGGGAGCTACCCTTACGATGTCCCGGATTACGCCTAATAGCGCATTGGAAGTGGATAACGGATCCGAATTCGAGCTCCGTCGACAAGCTTGCGGCCGCACTCGAGTGAGATCCGGCTGCTAACAAAGCCCGAAAGGAAGCTGAGTTGGCTGCTGCCACCGCTGAGCAATAACTAGCATAACC |
| VP19 | WKPEPCVY | Range of ML model scores | GCGAAATTAATACGACTCACTATAGGGGAATTGTGAGCGGATAACAATTCCCCTCTAGAAATAATTTTGTTTAACTTTAAGAAGGAGATATACATATGCTGGCCGAACTTAGCGAAGAAGCGTTGATCGAGGGGCGCTGGAAACCCGAACCCTGTGTCTACGGGTCTGGGAGCTACCCTTACGATGTCCCGGATTACGCCTAATAGCGCATTGGAAGTGGATAACGGATCCGAATTCGAGCTCCGTCGACAAGCTTGCGGCCGCACTCGAGTGAGATCCGGCTGCTAACAAAGCCCGAAAGGAAGCTGAGTTGGCTGCTGCCACCGCTGAGCAATAACTAGCATAACC |
| VP20 | FVAMCGAC | Multiple Cys in core | GCGAAATTAATACGACTCACTATAGGGGAATTGTGAGCGGATAACAATTCCCCTCTAGAAATAATTTTGTTTAACTTTAAGAAGGAGATATACATATGCTGGCCGAACTTAGCGAAGAAGCGTTGATCGAGGGGCGCTTTGTTGCTATGTGCGGAGCATGTGGGTCTGGGAGCTACCCTTACGATGTCCCGGATTACGCCTAATAGCGCATTGGAAGTGGATAACGGATCCGAATTCGAGCTCCGTCGACAAGCTTGCGGCCGCACTCGAGTGAGATCCGGCTGCTAACAAAGCCCGAAAGGAAGCTGAGTTGGCTGCTGCCACCGCTGAGCAATAACTAGCATAACC |
| VP21 | WGFASSCC | Multiple Cys in core | GCGAAATTAATACGACTCACTATAGGGGAATTGTGAGCGGATAACAATTCCCCTCTAGAAATAATTTTGTTTAACTTTAAGAAGGAGATATACATATGCTGGCCGAACTTAGCGAAGAAGCGTTGATCGAGGGGCGCTGGGGATTTGCATCCTCGTGCTGCGGGTCTGGGAGCTACCCTTACGATGTCCCGGATTACGCCTAATAGCGCATTGGAAGTGGATAACGGATCCGAATTCGAGCTCCGTCGACAAGCTTGCGGCCGCACTCGAGTGAGATCCGGCTGCTAACAAAGCCCGAAAGGAAGCTGAGTTGGCTGCTGCCACCGCTGAGCAATAACTAGCATAACC |

#### mRNA Library Creation

##### Library Assembly

An initial extension reaction (25µL) containing 1x Q5 Reaction buffer, 2.5 mM MgCl2, 0.25 mM dNTPs, 1 µM forward primer (P2), 1 µM reverse primer mix (P3-P10), 0.02 units/µL Q5 DNA Polymerase (NEB M0491L), and diluted in MQ-H2O. This was run for 5 cycles on a thermocycler at conditions listed in the extension reaction table (below). Once complete a 500 µL PCR amplification was set up containing all of the initial extension reaction, 1x Q5 Reaction buffer, 2.5mM MgCl2, 0.25mM dNTPs, 0.3µM forward primer (P1), 0.3µM reverse primer (P11), 0.02 units/µL Q5 DNA Polymerase (NEB M0491L), and diluted in MQ-H2O. This was carried out under conditions listed below in the PCR-2 amplification table. The progress of this reaction was checked by 3% agarose gel (supplemented with 1.5% ethidium bromide). Once sufficient amplification 1x volume of phenol/chloroform/isoamyl alcohol was added to the solution and centrifuged. The top layer was extracted, transferred to a clean tube, and 1x volume of chloroform/isoamyl alcohol was added to the solution and centrifuged. The top layer was again extracted, transferred to a clean tube, and then a 1/10 volume of 3M NaCl and 2x volume of 100% ethanol was added. This mixture was centrifuged for 10mins to pellet the DNA. Once pelleted, the DNA was washed with excess 70% ethanol, then allowed to dry at RT. The DNA was then solubilized in a 1/10 volume (50µL) of the PCR reaction.

Extension reaction conditions:

| Step | Temperature (°C) | Time (sec) |  |
| --- | --- | --- | --- |
| Initial Denature | 95 | 60 |  |
| Annealing | 50 | 60 | 5 cycles |
| Elongation | 72 | 60 |  |
| Final Elongation | 72 | 120 |  |

PCR-2 amplification conditions:

| Step | Temperature (°C) | Time (sec) |  |
| --- | --- | --- | --- |
| Initial Denature | 95 | 60 |  |
| Denature | 95 | 40 | 12 cycles |
| Annealing | 50 | 40 |  |
| Elongation | 72 | 40 |  |
| Final Elongation | 72 | 120 |  |

##### Transcription

A 500 µL transcription reaction (adapted from NEB T7 RNA polymerase protocols) containing amplified DNA (50 μL), 1x T7 RNA Polymerase Buffer, 4 mM Dithiothreitol (DTT), 16.5 mM MgCl2, 5 mM rNTPs, and 5 units/µL T7 RNA Polymerase (NEB M0251) was incubated overnight at 37 ºC. Upon successful transcription, magnesium pyrophosphate precipitates and the solution appears cloudy. Then, 57.5 µL of 10x Dnase I buffer and 15 µL of Dnase I (Promega PR-M6101) was added to the solution and incubated at 37 ºC for 1h. After DNAse incubation, EDTA, NaCl, and 100% isopropyl alcohol were added to final concentrations of 37.5 mM, 150 mM, 45%, respectively, upon which the solution became clear. The solution was then centrifuged at 4,000 rpm for 15 mins to pellet the RNA. The supernatant was removed, and the pellet was washed with excess 70% ethanol. This solution was briefly centrifuged at 4,000 rpm, the supernatant was removed, and the RNA pellet was allowed to dry at RT. The RNA pellet was then solubilized in 50 µL of MQ-H2O and an equal volume (50 µL) of 2x RNA loading dye. The sample was heated at 95 ºC for 2 mins, then run on a large scale 8% Urea-PAGE gel at 230 V for 1.5 h in a Tris-Borate-EDTA (TBE) buffer. The desired RNA band was then visualized under UV illumination at 254 nm on a silica-coated thin-layer chromatography plate and excised from the gel. The excised band was crushed into fine pieces and the RNA was extracted with 0.3 M NaCl (2x 1 hr incubation at RT). To collect the RNA, the gel was pelleted by centrifugation at 4,000 rpm and the supernatant isolated. The supernatant was then passed through a 0.45 µm filter before addition of 2x volume of 100% ethanol. The solution was mixed vigorously, then was centrifuged at 4,000 rpm for 15 mins to pellet the RNA. The pellet was washed with excess 70% ethanol, briefly centrifuged at 4,000 rpm, and then allowed to dry at RT. The RNA was solubilized in MQ-H2O and the concentration was determined by nanodrop. RNA was stored at -20 ºC until use.

##### Puromycin Ligation

RNA for each library was covalently linked to puromycin via an adapted Y-ligation strategy. The reaction was carried out in MQ-H2O with 1µM RNA, 20% DMSO, 1.5µM P-Linker (d(pCTCCCGCCCCCCGTCC)-(SPC18)5-d(CC)-puromycin), 1x T4 RNA ligase buffer, and 1unit/µL T4 RNA ligase I (PR-M1051). Once assembled the reaction was incubated at 37ºC for 0.5h. After incubation 1x volume of crashout solution (0.6M NaCl, and 50mM EDTA), 0.03x volume of 100% glycogen, and 2.2x volume 100% ethanol were added. This mixture was briefly vortexed and then centrifuged at 15,000xg for 10mins to pellet the Plinked RNA. The supernatant was removed and excess 70% ethanol was added, briefly centrifuged at 15,000xg, supernatant removed, and the pellet was dried at RT. Once dried the Plinked RNA pellet was reconstituted in an equal volume as the input RNA in MQ-H2O. The efficiency of the reaction was determined by 8% Urea Gel (National Diagnostics, EC-833).

#### LynD Substrate Tolerance mRNA Display Assay Development

##### Translation and Reverse Transcription

Translation of puromycin-linked RNA was performed using a custom NEB PURExpress® kit Δ(AA, tRNA, -RF123, NEB: E6840S). Library L1 was translated without restriction factors at 10 μL total translation volume (split into two 5 µL replicates) for each input sample. Translations were incubated at 37 ºC for 30 mins, followed by a 10 min incubation at RT to facilitate fusion of peptide to its mRNA strand. EDTA was then added to a final concentration of 17 mM to dissociate the ribosome, and the mixture was incubated at 37 ºC for 30 mins. Then, complementary DNA was appended by a reverse transcription reaction containing all the translation product, 0.625 mM dNTPs, 5 μM reverse primer (P11), 62.5 mM Tris-HCl pH 8.3, 37.5 mM Mg(OAc)_2_, 25 mM KOH, 2.5 x M-MLV reverse transcriptase H (-) point mutant (Promega, M3681, supplied at 40x), and MQ-H2O (to reach final volume). This solution was incubated at 42 ºC for 1h and a 0.5 µL sample was taken for future qPCR analysis.

##### LynD Treatment and Free Cysteine Biotinylation

After reverse transcription, the library was treated with LynD to form thiazoles in a reaction containing the entire reverse transcription solution, 50 mM HEPES pH 7.5, 75 mM NaCl, 0.5 mM TCEP, 5 mM MgCl_2_, 5 mM ATP, 10.3 µM LynD, and MQ-H_2_O (to reach final volume). The thiazole-forming reaction was run at room temperature for 30 min and a 0.5 µL sample was taken for future qPCR analysis. Then TCEP was added to a final concentration of 2.5 mM in TBS buffer (25 mM Tris-HCl pH 7.5, 150 mM NaCl). The mixture was incubated on ice for 30 min, then incubated with 1x biotin iodoacetamide (BIAA) on ice for 1 hr.

##### Purification

The library was then HA purified. Anti-HA magnetic beads (Thermo Fischer Scientific, 88836) were prepared at a 4:1 ratio of bead slurry to initial IVT volume, washed 3x with TBS-T (25 mM Tris-HCl pH 7.5, 150 mM NaCl, 0.05% Tween-20), and resuspended in blocking buffer (TBS-T, 2 mg/mL BSA, 2 mg/mL yeast RNA) to a final volume of 50 µL when mixed with each library sample. The library-HA bead solutions were mixed at room temperature for 30 min, washed 3x with TBS-T, then eluted in 50 µL of a 1:1 ratio of TBS-T to HA synthetic peptide (Thermo Fischer Scientific, 26184) by mixing at room temperature for 1 hr. Each sample was mixed with 3 µL glycogen and a 3x volume of ice-cold acetone and incubated at -20°C for 10 min. The library was recovered by centrifugation at 15,000 g for 15 min. The supernatant was discarded and the samples were air-dried at room temperature for 5 min. The dried library samples could then be stored at -20°C until further use.

##### Streptavidin Pulldown

The purified, dry library samples were resuspended in 17.5 µL blocking buffer and a 5 µL sample was taken for future qPCR analysis. The remaining solution was then split into two 6 µL replicates. SA magnetic beads (Thermo Fisher Scientific, 11205D) were prepared at a 4:1 ratio of bead slurry to initial IVT volume, washed 2x with TBS-T, washed 1x with blocking buffer, and resuspended with blocking buffer to a final volume of 50 µL when mixed with each library sample. The library-SA bead solutions were mixed at room temperature for 30 min (the supernatant was kept), washed 2x with Urea in TBS-T (1 M Urea, 25 mM Tris-HCl pH 7.5, 150 mM NaCl, 0.05% Tween-20) for 5 min, and washed 2x with TBS-T. The SA beads were then dropped in 100 µL TBS-T, hand-transferred to 50 µL 1x Q5 buffer, and then heated at 95 °C for 5 min to elute the cDNA of any SA bead-bound library members. The 5 µL of the initially kept supernatant was diluted 10-fold in 1x Q5 buffer.

##### qPCR for Assay Validation

An Applied Biosystems (AB) ViiA 7 RealTime PCR System (qPCR) was used for amplification and analysis after mRNA display experiments. Each sample was analyzed in a 10µL solution of 1x SsoAdvanced Universal SYBR Green Supermix (Bio-Rad, 172-5271), 0.125 µM Forward and Reverse primers (P1, P11), 2µL of respective sample, and diluted to the final volume with MQ-H2O. qPCR standards were prepared by reverse transcription of a known quantity of RNA into cDNA, assumed 100% yield, and dilutions to 2e9 , 2e8 , 2e7 , 2e6 , 2e5 , 2e4 molecules were carried out. The qPCR method was carried out by first heating at 50ºC for 2 mins followed by 10 min 95ºC incubation. Cycling was then carried out between 95ºC for 15 sec and 60ºC for 1 min, with cycler heating acceleration held at 1.6ºC between each step. A total of 40 cycles was carried out in each qPCR run. During each elongation step, SYBR green fluorescence was measured with ROX as a passive reference. Following the run, standard curves were generated and used to calculate cDNA quantities.

##### PCR Amplification and Preparation for NGS

The cDNA of the purified library input, 10-fold diluted SA supernatant, and SA elution samples were amplified according to standard Q5 polymerase amplification protocols. All PCR reactions were performed alongside a no template control. Amplification was performed until bands could be observed via DNA gel, usually 15-30 cycles.

PCR amplification for NGS conditions:

| Step | Temperature (°C) | Time (sec) |  |
| --- | --- | --- | --- |
| Initial Denature | 95 | 60 |  |
| Denature | 95 | 40 | 15-30 cycles |
| Annealing | 69 | 40 |  |
| Elongation | 72 | 40 |  |
| Final Elongation | 72 | 120 |  |

##### NGS Sample Preparation

Recovered DNA from the end of the mRNA display round was prepared for Azenta Amplicon EZ NGS analysis. A standard Q5 polymerase reaction was carried out for 12 cycles (using forward primer P12 and one of the reverse primers P13-16). After sufficient amplification was observed the sample was cleaned up using a standard column purification kit and the concentration determined by nanodrop. Then 500µg were sent (in MQ-H2O) for NGS.

#### NGS Data Processing for Substrate Tolerance Information

##### Initial NGS Data Cleanup

Raw NGS data were processed by collecting valid open reading frames (DNA sequences) of the expected length, which were then in silico translated into peptide sequences. When available, forward and reverse NGS results were combined into one dataset. Prior to modeling and analysis, peptide sequences were curated by extracting the variable region, combining duplicate reads, and removing any peptides appearing in both selection and antiselection datasets. The final curated round 3 dataset was nearly balanced, with 469,867 selection and 491,952 antiselection peptides.

##### Y* score and S-Score Calculations

The S-score is a measure of the expected fitness of a given peptide in the absence of any epistatic effects. To calculate S-scores, we first calculate the Y* score matrix for a given dataset following the method described in Vinogradov et al.^1^ Briefly, this involves calculating the ratio of selection vs antiselection reads containing each amino acid at each position, with a higher value indicating more favorable residue/position combinations. We then transform these values onto a log_2_ scale, as is commonly done for interpretability. We calculate Y* scores for specific sublibraries by selecting only peptides with a Cys residue at the target position (e.g., C6). To calculate the S-score of a given peptide, we simply sum the Y* score contribution for each individual residue/position coordinate across the variable region. To evaluate the predictive power of the S-score, we calculated the Y* score matrix on a random 90% split of the dataset and inferred S-scores for the remaining 10% of peptides. A Gaussian mixture model was used to fit the S-score distribution using either one or two Gaussian components to determine μ and σ values.

##### Machine Learning Model Creation

To learn the substrate tolerance patterns directly from the selection data, we trained a simple multilayer perceptron (MLP) to classify peptides as ‘modified’ (1) or ‘unmodified’ (0). The curated experimental data was randomly split into training/validation/test sets using an 80/10/10 split. Peptides variable region sequences were one-hot encoded to obtain 8 x 20 = 160 input features. The MLP included a single hidden layer of 128 dimensions with 10% dropout and a ReLU activation function, and the final classification output was transformed with a Sigmoid function. Training used a batch size of 2048 and a learning rate of 10-3 with the AdamW optimizer for up to 50 epochs. The best model checkpoint was selected according to max validation set accuracy. Trained model checkpoints as well as code for training, validation, and data analysis are provided on GitHub at <https://github.com/Kuhlman-Lab/LynD-substrate-modeling>.

##### Model Evaluation

The primary metrics used for model evaluation were balanced accuracy (BA, **Figure S3**) and positive predictive value (PPV). To compare S-score and MLP model predictions on a common scale (0-1), we transformed S-scores into S-score percentiles before analysis. To calculate epistasis between positions using our MLP model, we adapted the protocol described by Vinogradov et al. to calculate *epi* scores.^1^ To generate the +1/-1 position map, we averaged the *epi* scores calculated individually for the +1/-1 positions of sublibraries 5-8. We quantified the contributions of each residue to overall peptide fitness for validation peptide LSMQKCYS by fixing individual residues and allowing the remainder (marked by ‘X’) to vary randomly. Each partially random library was evaluated *in silico* by sampling 10^6^ peptides. To evaluate the fraction of the theoretical complete library modifiable by LynD, we generated 10^7^ random peptides with at least one Cys residue and calculated the fraction of peptides predicted to have >90% modification by our MLP model.

#### Validation Peptide Experimental Design

##### Validation Peptide Nomination

Peptides were generated for nomination as follows. To assess the effect of the cysteine position on modification efficiency (**Genes 1-6**), a set of peptides with a sliding cysteine motif was designed. These genes used the conserved motif **ACM** and filled the remaining positions with alternating glycine-serine repeats on each side. To collect peptides with a wide range of predicted modification scores (**Genes 7-19**), we generated 10^6^ random peptides with a Cys residue at variable position 6. Any peptides highly similar (within a Hamming distance of 2) to any peptide in the training set were removed to avoid data leakage. The remaining peptides were then scored by S-score and MLP models, and validation peptides were selected manually to represent a range of predicted modification and concurrence between the two scoring methods. To demonstrate that our model could identify highly modifiable peptides with two Cys residues (**Genes 20** and **21**), we generated a library of 10^6^ random peptides with exactly two Cys residues. We then filtered them by Hamming distance, as described above, and by predicted modification efficacy (>0.97), followed by manual selection.

##### Gene Preparation

Validation peptide gene fragments for IVT were first resuspended to a final concentration of 10 ng/μL. PCR amplification was carried out under standard Q5 DNA polymerase (NEB M0491) protocols. P17 and P18 were used to amplify gene fragments for IVT. Amplification was performed under conditions listed in the below table and confirmed by DNA gel (3% agarose gel supplemented with 1.5% ethidium bromide). DNA was then purified by a PCR purification kit, isolated in MQ-H2O, and concentration was determined by nanodrop. Gene fragments were stored at -20 °C until further use.

Gene fragment PCR conditions:

| Step | Temperature (°C) | Time (sec) |  |
| --- | --- | --- | --- |
| Initial Denature | 95 | 120 |  |
| Denature | 95 | 30 | 25 cycles |
| Annealing | ** | 30 |  |
| Elongation | 72 | 120 |  |
| Final Elongation | 72 | 300 |  |

**Primer annealing temperatures were determined by NEB Tm calculator

##### IVT and LynD Treatment

All IVT assays were carried out using PURExpress® Δ (aa, tRNA) Kit (NEB E6840S) on a 5 uL scale. The solution was first incubated at 37ºC for 0.5h to facilitate translation. A 2.5 µL sample was removed for future analysis as the IVT input. The rest of the sample was treated with LynD to form thiazoles with 50 mM HEPES pH 7.5, 75 mM NaCl, 0.5 mM TCEP, 5 mM MgCl_2_, 5 mM ATP, 10.3 µM LynD, and MQ-H_2_O (to reach final volume). The thiazole-forming reaction was run at room temperature for 30 min.

##### Desalting Protocol and Preparation for Mass Spectrometry

The peptide was purified from the reaction solutions using a C18 Spin tip (Pierce PI84850). To prepare the C18 spin tip, 15 uL of a C18 elution solution (80% acetonitrile and 0.5% acetic acid in water) was added, spun down, and flow through removed. Then 15 uL of a C18 elution solution (4% acetonitrile and 0.5% acetic acid in water) was added, spun down, and flow through removed. Next the reaction sample solution was added and washed twice with the wash solution. Finally, 1.5uL of a half-saturated solution of α-cyano-4-hydroxycinnamic acid in elution buffer was used to elute the peptide from the C18 tip. This was spotted on a MALDI-TOF plate and analysis was carried out on an Applied Biosystems AB SIEX TOF/TOF 5800 system in reflector positive mode (**Figure S4**).

#### LynD Cloning, Expression, and Purification.

The genes for LynD was purchased as gene blocks from Twist Biosciences, PCR amplified, and cloned into pMCSG7 vectors using the following primers:

LynD-Forward (5’→3’): TACTTCCAATCCAATGCGATGCAAAGCACACCACTGC

LynD-Reverse (5’→3’): TTATCCACTTCCAATGCGCTATTAGAACGGCATTGGGGTC

The PCR product was phosphorylated with T4 PNK and treated with T4 DNA polymerase to create LIC-overhangs. Plasmid pMCSG7 were linearized with SSPI, dephosphorylated with Antarctic Phosphatase, and treated with T4 DNA Polymerase to create LIC-overhangs. Prepared vectors and PCR products were incubated for 10 min at 25°C and then transformed into One-Shot® Top 10. pMCSG7-LynD was transformed and expressed in BL21-CodonPlus (DE3)-RIPL Competent Cells.

His6-LynD protein was expressed and purified from E. coli RIPL cells as in the following report.^2^ A 5 mL saturated culture was used to inoculate 1 L of Luria-Bertani (LB) medium supplemented with ampicillin (100 μg/mL) and chloramphenicol (34 μg/mL). The culture was incubated at 37°C and shaken at 220 rpm until reaching an OD600 of 0.6-0.8 at which point the medium was supplemented with 0.2 mM IPTG. Culture was then cooled to 18°C and grown overnight (~20 hours). Cells were pelleted and stored at -80°C until purification. Pellets were resuspended in 40 mL of wash buffer (20 mM Tris pH 8.0, 200 mM NaCl, 50 mM Imidazole, and 1 mM TCEP) supplemented with 1.5 mM Phenylmethylsulfonyl fluoride (PMSF), 80 units of DNAseI, 0.25 mg lysozyme and sonicated twice with intermittent pulses with a 30% maximum amplitude for 1:30 min. The soluble protein was recovered by pelleting the cell debris by centrifugation at 4°C 15,000 rpm for 60 min and was filtered through a 0.45 μm sterile syringe filter. The supernatant was loaded onto a 5-mL HisTrap (Ni2+) IMAC column and washed with 5 column volumes (CV) of wash buffer. Protein was eluted with elution buffer (20 mM Tris pH 8.0, 200 mM NaCl, 500 mM imidazole, 1 mM TCEP) over a gradient 0-100% over 10 CV and checked by SDS-PAGE (**Figure S7**). Fractions collected which contained purified protein were collected and concentrated to 2.5 mL using a Centricon (10,000 Da MWCO) concentrator (EMD Millipore®). The concentrated protein was then buffer exchanged using PD-10 column (GE Healthcare Life Sciences®) into storage buffer (10 mM HEPES pH 7.4, 150 mM NaCl, 1 mM TCEP and stored at -80°C. A 1 L culture yielded ~22 mg of protein.

His6-LynD Sequence: MHHHHHHSSGVDLGTENLYFQSNAMQSTPLLQIQPHFHVEVIEPKQVYLLGEQANHALTGQLYCQILPLLNGQYTLEQIVEKLDGEVPPEYIDYVLERLAEKGYLTEAAPELSSEVAAFWSELGIAPPVAAEALRQPVTLTPVGNISEVTVAALTTALRDIGISVQTPTEAGSPTALNVVLTDDYLQPELAKINKQALESQQTWLLVKPVGSVLWLGPVFVPGKTGCWDCLAHRLRGNREVEASVLRQKQAQQQRNGQSGSVIGCLPTARATLPSTLQTGLQFAATEIAKWIVKYHVNATAPGTVFFPTLDGKIITLNHSILDLKSHILIKRSQCPTCGDPKILQHRGFEPLKLESRPKQFTSDGGHRGTTPEQTVQKYQHLISPVTGVVTELVRITDPANPLVHTYRAGHSFGSATSLRGLRNTLKHKSSGKGKTDSQSKASGLCEAVERYSGIFQGDEPRKRATLAELGDLAIHPEQCLCFSDGQYANRETLNEQATVAHDWIPQRFDASQAIEWTPVWSLTEQTHKYLPTALCYYHYPLPPEHRFARGDSNGNAAGNTLEEAILQGFMELVERDGVALWWYNRLRRPAVDLGSFNEPYFVQLQQFYRENDRDLWVLDLTADLGIPAFAGVSNRKTGSSERLILGFGAHLDPTIAILRAVTEVNQIGLELDKVPDENLKSDATDWLITEKLADHPYLLPDTTQPLKTAQDYPKRWSDDIYTDVMTCVNIAQQAGLETLVIDQTRPDIGLNVVKVTVPGMRHFWSRFGEGRLYDVPVKLGWLDEPLTEAQMNPTPMPF

#### Electrostatic Calculations

We used the Rosetta FlexPepDock protocol^3^ to dock the native LynD substrate for structural analysis (PDB ID: 4V1T). We used AlphaFold3^4^ to obtain a computational model of LazE, which we aligned onto the docked LynD/peptide complex for comparison. The APBS web server^5^ was used to calculate full electrostatic potential grids using the AMBER force field, and the docked peptide was used as a reference point to sample the neighborhood around each residue. A square 5Å grid around the Cβ of each residue was sampled to obtain the mean electrostatic potential using the GridDataFormats python package (**Figure S9**).

### Supplemental Figures

| **A** | **B** |
| --- | --- |
| \| **Average cDNA count**; (+) LynD Input \| 3.67E+08; (-) LynD Input \| 6.56E+08; (+) LynD SA Unbound \| 3.17E+08; (-) LynD SA Unbound \| 7.88E+07; (+) LynD SA Bound \| 7.02E+07; (-) LynD SA Bound \| 4.18E+08 |  |

|  | Percent cDNA Recovery |
| --- | --- |
| (+) LynD SA Unbound | 86.3324 |
| (+) LynD SA Bound | 19.1275 |
| (-) LynD SA Unbound | 12.021 |
| (-) LynD SA Bound | 63.7108 |

Figure S1. Streptavidin capture assay validation using test gene G1. (A) cDNA counts quantized by qPCR. (B) Percent cDNA recovery with and without LynD. Percent cDNA recovery were calculated by dividing the SA bound or unbound fraction by the cDNA counts after the acetone purification input sample. Error bars are one standard deviation, calculated from two replicates.

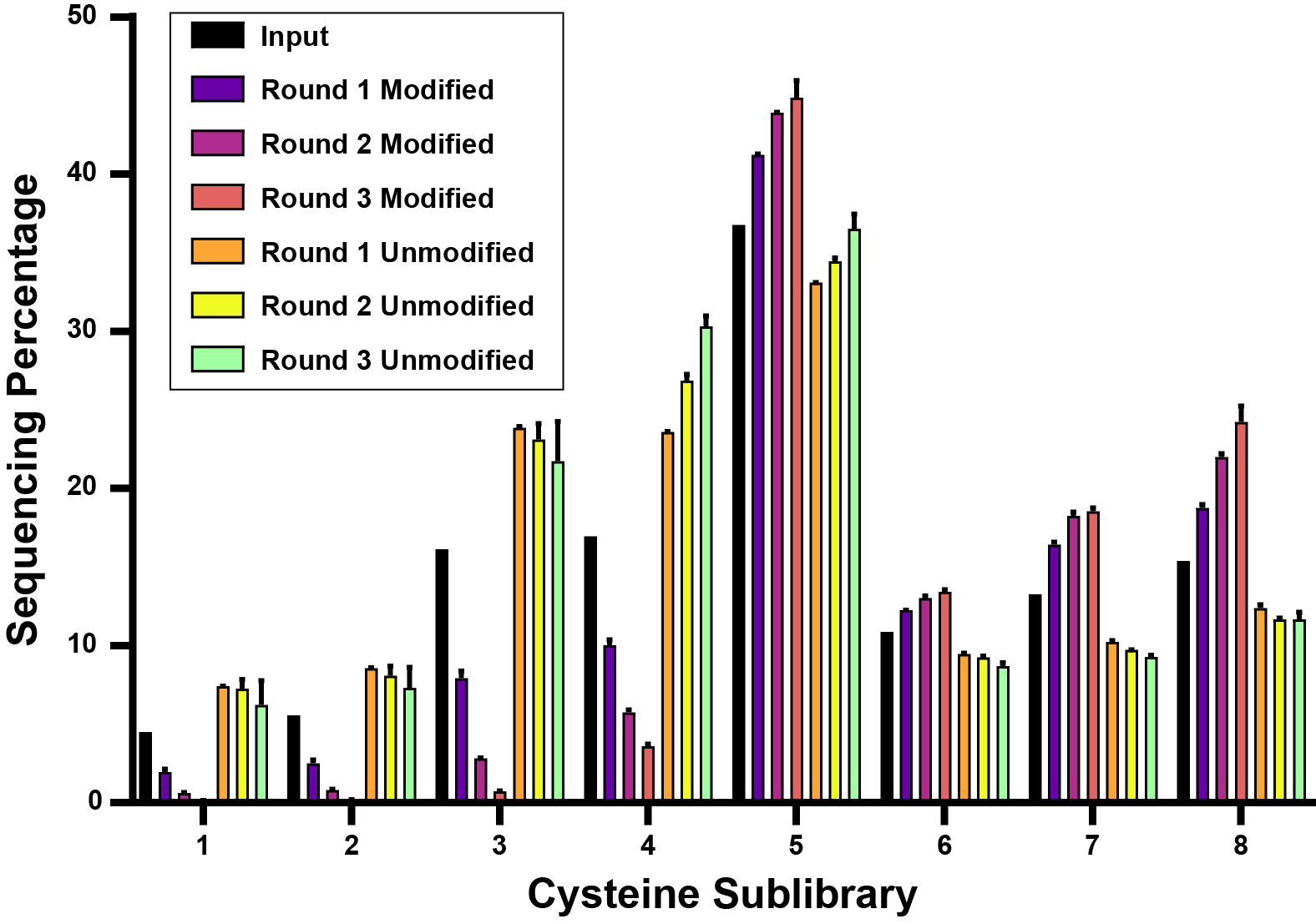

Figure S2. Sublibrary sequencing distribution over three rounds. Cysteine sublibraries are separated by fixed cysteine position.

| ***Round*** | ***Modified Peptides*** | ***Unmodified Peptides*** | ***S-score***  ***Balanced Accuracy*** | ***MLP***  ***Balanced Accuracy*** |
| --- | --- | --- | --- | --- |
| *1* | *440058* | *620180* | *67.5%* | *70.7%* |
| *2* | *210392* | *331083* | *79.5%* | *84.5%* |
| *3* | *469867* | *491952* | *84.6%* | *91.5%* |

Figure S3. Balanced accuracy comparison between S-Score and MLP models over three rounds.

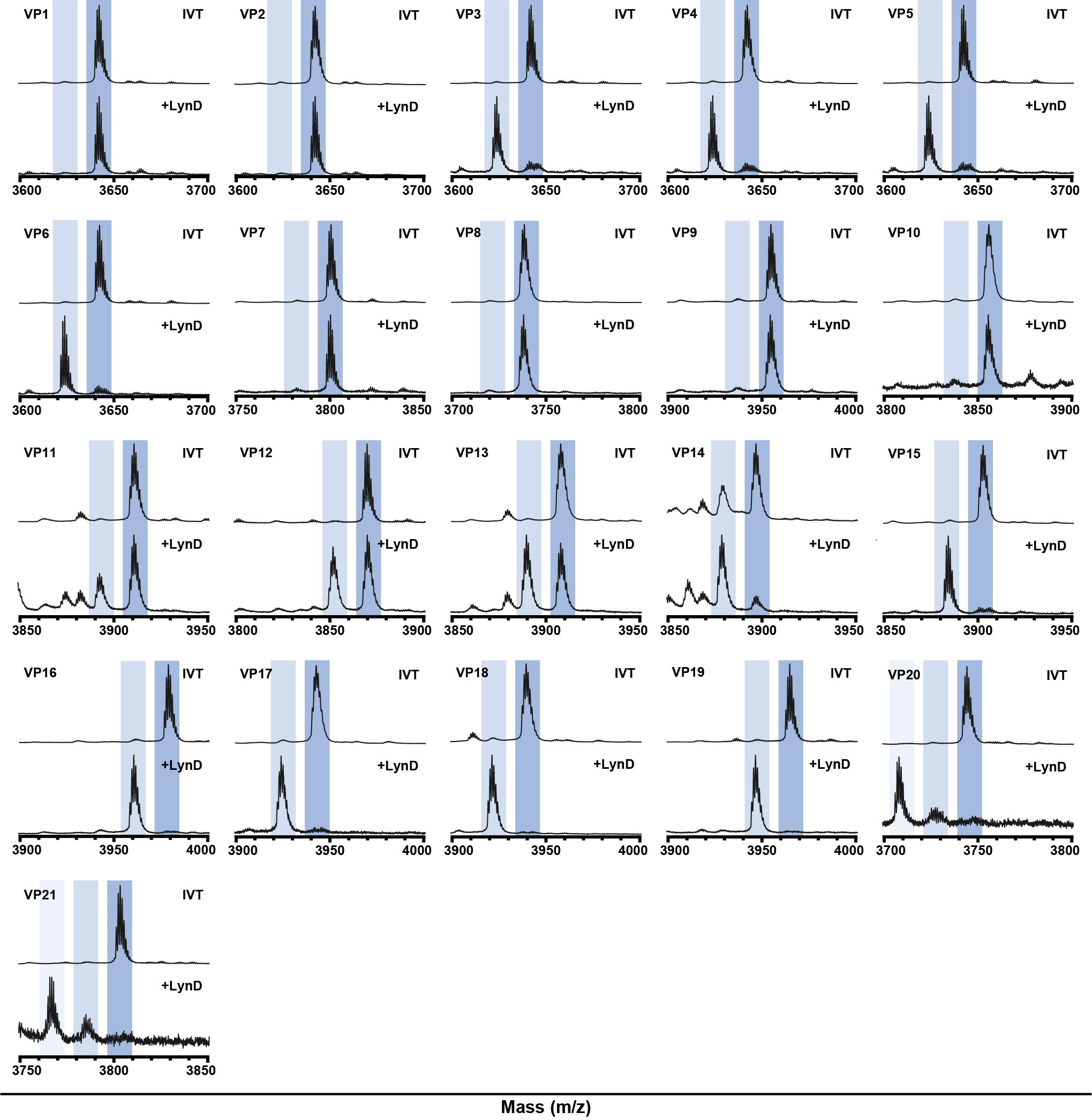

Figure S4. Mass spectrometry data for validation peptides before and after LynD treatment. Blue bars represent expected masses for the cysteine-containing peptide substrates, with lighter blues indicating the relative masses for thiazoline formations.

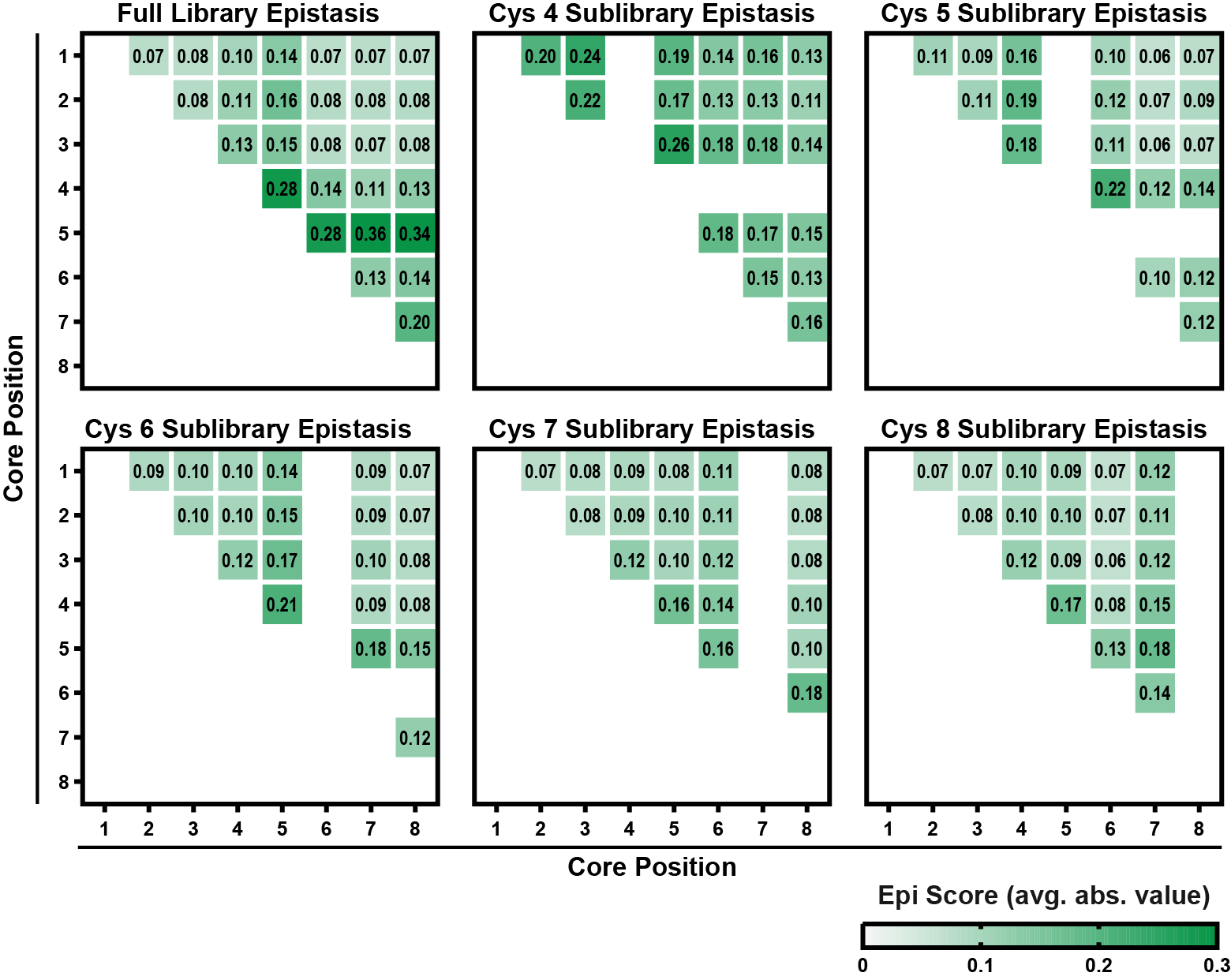

Figure S5. Positional epistasis plots for sublibraries 4-8. Sublibraries are separated by fixed cysteine position.

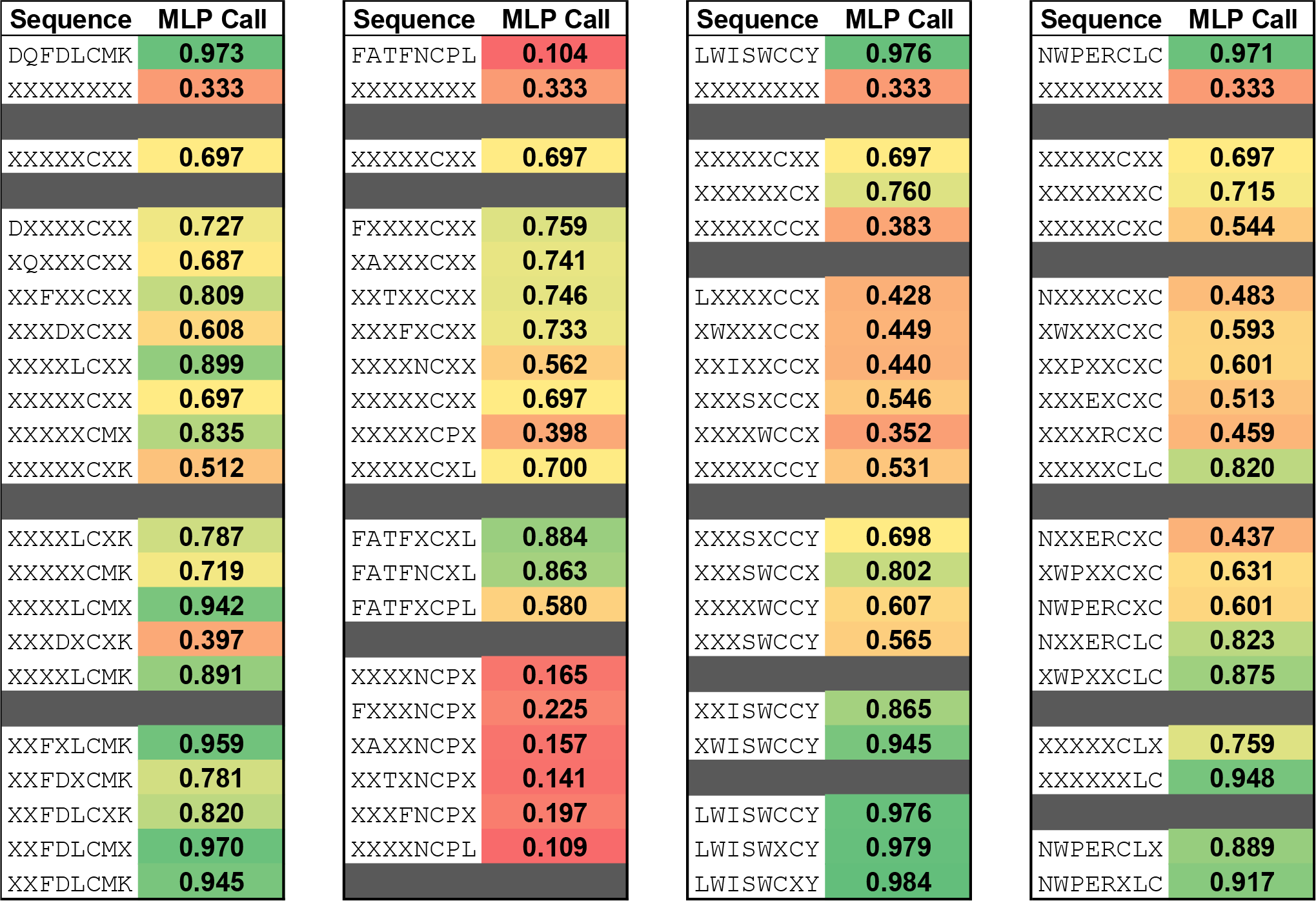

Figure S6. Epistasis interactions breakdown for example peptide substrates. Randomized core positions are represented with an X. The MLP call is the average MLP model predicted modification efficiency.

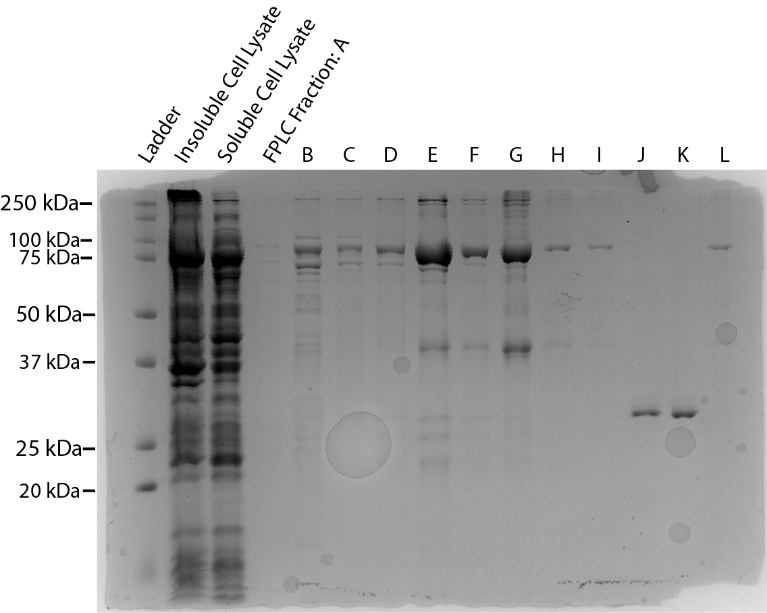

Figure S7. SDS-PAGE gel of LynD FPLC purification fractions. FPLC fractions D-F were collected as the purest material.

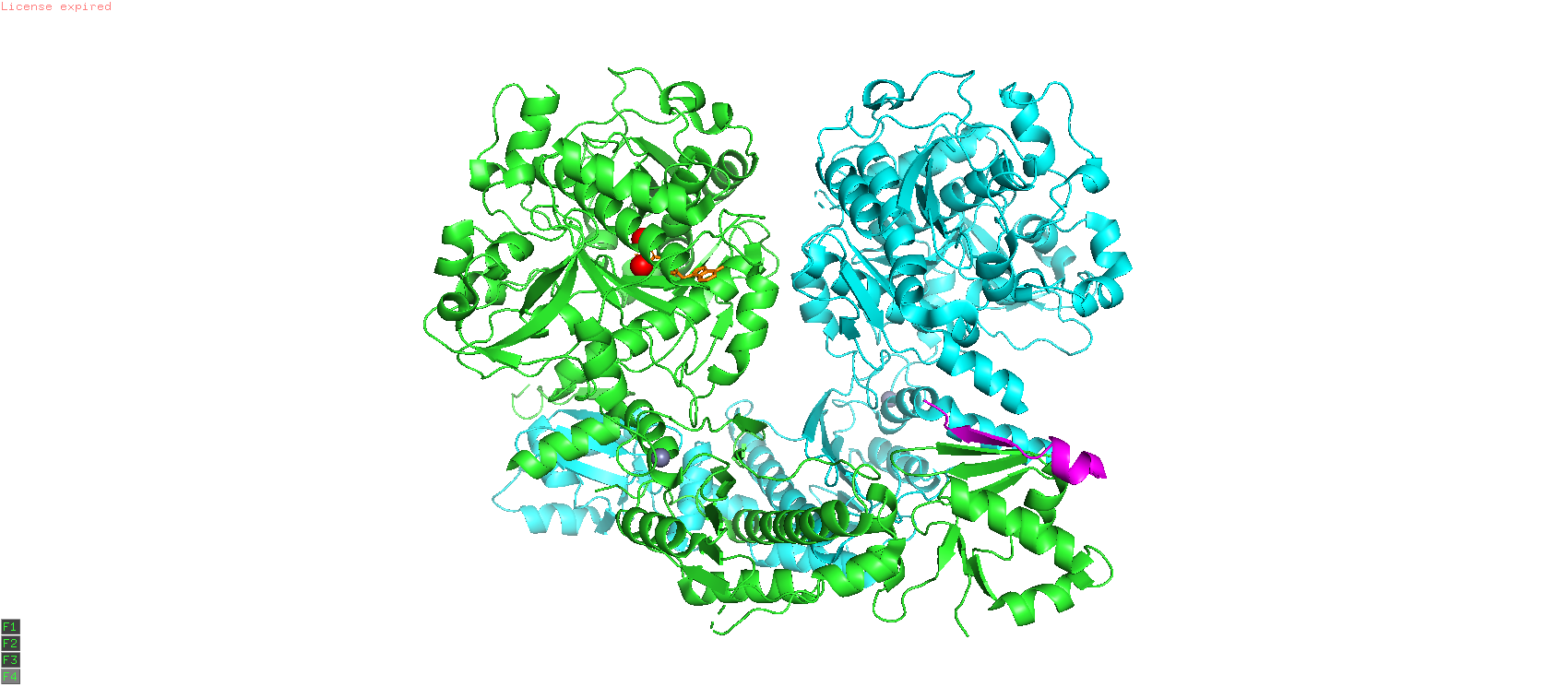

Figure S8. LynD homodimer crystal structure. Green and blue monomers are depicted with the substrate leader peptide (magenta) bound. Magnesium (red) and ATP (orange) are also shown bound to the green LynD monomer. PDB: 4V1V.

**A**

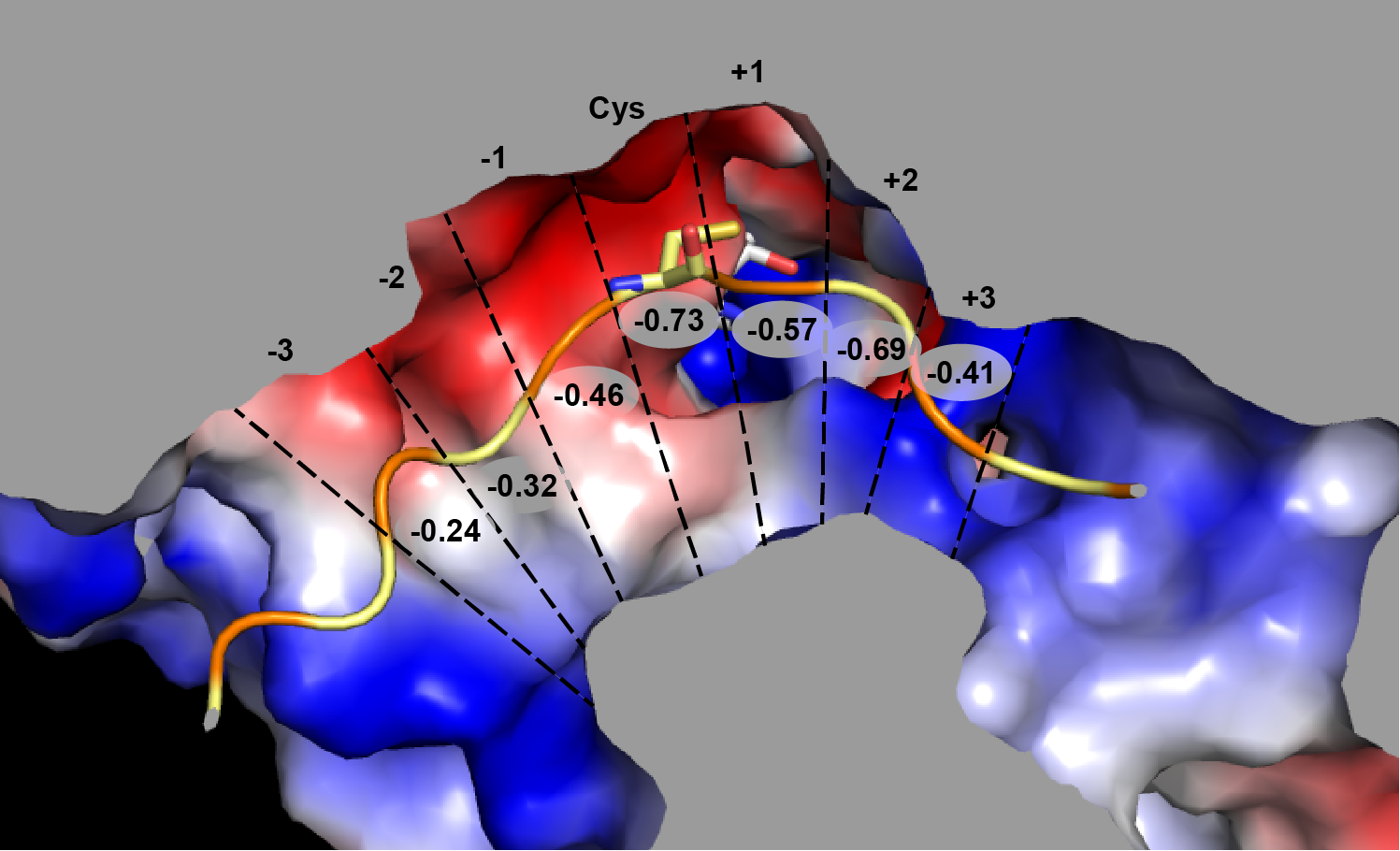

B

| **Residue position relative to Cys** | **-3** | **-2** | **-1** | **0** | **1** | **2** | **3** |
| --- | --- | --- | --- | --- | --- | --- | --- |
| **LynD (mV)** | -0.24 | -0.32 | -0.46 | -0.73 | -0.57 | -0.69 | -0.41 |
| **LazE (mV)** | -0.20 | -0.47 | -0.78 | -1.58 | -2.18 | -1.15 | -0.80 |
| **LynD – LazE (mV)** | -0.04 | +0.15 | +0.32 | +0.85 | +1.61 | +0.46 | +0.39 |

Figure S9. (A) Electrostatic potential map of LynD catalytic site with AMP (white) and the docked peptide substrate (alternating orange and yellow for each residue). The substrate cysteine stick representation is shown. The electrostatic potential map, as generated by PyMol, compares negative regions (red) to more positive regions (blue) and is sliced (grey) to distinguish the catalytic pocket. Circled numbers represent the calculated mean electrostatic potential for each docked residue. Note that dashed lines are a graphical representation and do not mark the boundaries for the electrostatic potential grids around each residue. (B) Mean electrostatic potential calculations for LynD and LazE. Calculations were made for three residues upstream and downstream of the docked cysteine in the catalytic site as described.
